# The G protein biased serotonin 5-HT 2A receptor agonist lisuride exerts anti-depressant drug-like activities in mice

**DOI:** 10.1101/2023.06.01.543310

**Authors:** Vladimir M. Pogorelov, Ramona M. Rodriguiz, Bryan L. Roth, William C. Wetsel

## Abstract

There is now evidence from multiple Phase II clinical trials that psychedelic drugs can exert long-lasting anxiolytic, anti-depressant, and anti-drug abuse (nicotine and ethanol) effects in patients. Despite these benefits, the hallucinogenic actions of these drugs at the serotonin 2A receptor (5-HT2AR) limit their clinical use in diverse settings. Activation of the 5-HT2AR can stimulate both G protein and β-arrestin (βArr) -mediated signaling. Lisuride is a G protein biased agonist at the 5-HT2AR and, unlike the structurally-related LSD, the drug does not typically produce hallucinations in normal subjects at routine doses. Here, we examined behavioral responses to lisuride, in wild-type (WT), βArr1-KO, and βArr2-KO mice. In the open field, lisuride reduced locomotor and rearing activities, but produced a U-shaped function for stereotypies in both βArr lines of mice. Locomotion was decreased overall in βArr1-KOs and βArr2-KOs, relative to WT controls. Incidences of head twitches and retrograde walking to lisuride were low in all genotypes. Grooming was depressed in βArr1 mice, but was increased then decreased in βArr2 animals with lisuride. Prepulse inhibition (PPI) was unaffected in βArr2 mice, whereas 0.5 mg/kg lisuride disrupted PPI in βArr1 animals. The 5-HT2AR antagonist MDL100907 failed to restore PPI in βArr1 mice, whereas the dopamine D2/D3 antagonist raclopride normalized PPI in WTs but not in βArr1-KOs. Using vesicular monoamine transporter 2 mice, lisuride reduced immobility times in tail suspension and promoted a preference for sucrose that lasted up to 2 days. Together, it appears βArr1 and βArr2 play minor roles in lisuride’s actions on many behaviors, while this drug exerts anti-depressant drug-like responses without hallucinogenic-like activities.

## 1. Introduction

Lisuride was first synthesized in 1960 as an analog of methysergide (Zikán and Semonský, 1960) and, as an ergoline derivative, it has a chemical structure similar to that of *D*-lysergic acid diethylamide (LSD). Both lisuride and LSD bind with high affinities to serotonin (5-HT) 2A receptors (5-HT2AR) and signal through Gα_q_ by activation of phospholipase C leading to production of inositol phosphates and diacylglycerol with the release of intracellular Ca^2+^ *in vitro* (Hoyer et al., 1994; Egan et al., 1998; Kurrasch-Orbaugh et al., 2003; Cussac et al., 2008). Additionally, there is evidence that 5-HT2AR agonists can stimulate phospholipase A_2_ and mediate release of arachidonic acid (Felder et al., 1990; Kurrasch-Orbaugh et al., 2003). While lisuride and LSD are partial agonists at the 5-HT2AR (Egan et al., 1998; Kurrasch-Orbaugh et al., 2003; Wacker et al., 2013; Zhang et al., 2022), they bind also dopaminergic, adrenergic, and other serotonergic receptors (Piercey et al., 1996; Egan et al., 1998; Marona-Lewicka et al., 2002; Millan et al., 2002; Kroeze et al., 2015). Despite these similarities, LSD possesses hallucinogenic activity at doses as low as 20 µg (Greiner et al., 1958), while lisuride is devoid of these psychedelic effects when tested up to 600 µg in humans (Herrmann et al., 1977).

Although lisuride is reported to produce hallucinations in some patients with Parkinson’s disease (Schachter et al., 1979; Parkes et al., 1981; LeWitt et al., 1982; Vaamonde et al., 1991), these effects may be attributable to dysregulation of dopaminergic and other neurotransmitter systems in this disease. Nevertheless, the lack of hallucinations with lisuride in normal subjects seems surprising since agonism at the 5-HT2AR is known to mediate the hallucinogenic actions of psychedelics (Schreiber et al., 1994; Glennon et al., 1983; Keiser et al., 2009) and because antagonistic actions by atypical antipsychotic drugs diminishes the psychosis associated both with schizophrenia and Parkinson’s psychosis (Roth et al., 2004; Meltzer and Roth, 2013). To date, lisuride has been used in humans to treat migraine and cluster headache, parkinsonism, and hyperprolactinemia (Herrmann et al., 1977; Verde et al., 1980; McDonald and Horowski, 1983; Raffaelli et al., 1983). By comparison, LSD has potential efficacy in treating cluster headache, anxiety and depression in life-threatening situations when combined with psychotherapy, and it may be useful in studying consciousness and treating substance abuse (Sewell et al., 2006; Gasser et al., 2015; Bogenschutz et al., 2016; Carhart-Harris et al., 2016a,b; Griffiths et al., 2016; Davis et al., 2021; Goodwin et al., 2022, 2023; von Rotz et al., 2022). Hence, the therapeutic profiles of these drugs are rather different.

In rodents the head twitch response (HTR) to psychedelics is known to be mediated by 5-HT2AR activation (Glennon et al., 1984; Keiser et al., 2009; González-Maeso et al., 2007) and is used frequently to determine the potential psychedelic actions of drugs. However, it should be emphasized that several non-hallucinogenic drugs including quipazine, 5-hydroxytryptophan, ergometrine, cannabinoid CB1 antagonists, rolipram, fenfluramine, and other drugs also induce HTRs (Corne and Pickering, 1967; Malick et al., 1977; Wachtel, 1983; Darmani, 1998; Darmani and Pandya, 2000), although to our knowledge there are no classical psychedelic drugs that do not induce HTRs. Thus, while there are many false positives for the HTR, we are unaware of false negative hallucinogens. In contrast to LSD and other psychedelics, lisuride does not produce HTRs in rats (Gerber et al., 1985) or mice (González-Maeso et al., 2007; Halberstadt and Geyer, 2013). Besides HTRs, another distinction between lisuride and LSD is their differential agonist profile at the 5-HT2AR. While both ligands bind this receptor, the residues in the binding pocket bound by these ligands produce slightly different receptor conformations and may show differential responses through G protein and β-arrestin (βArr) mediated signaling (Kim et al., 2020; Cao et al., 2022). Parenthetically, the ability of ligands to stimulate or inhibit different pathways from the same receptor is termed functional selectivity (Urban et al., 2007) or biased signaling (Roth and Chuang 1987; Kenakin, 1995). Recently, LSD has been shown to be βArr biased at the 5-HT2AR *in vitro* (Wacker et al., 2017; Cao et al., 2022) and *in vivo* (Rodriguiz et al., 2021).

In present study we used the βArr1 and βArr2 mice to determine whether lisuride could exert differential effects on motor performance, PPI, and various ethological behaviors—including HTRs in the presence and absence of βArrs. As there is emerging evidence that psychedelics and biased 5-HT2AR agonists have long-lasting effects on depression (Carhart-Harris et al., 2016a,b; Griffiths et al., 2016; Davis et al., 2021; Goodwin et al., 2022, 2023; McClure-Begley and Roth, 2022; von Rotz et al., 2022), we tested if lisuride had anti-depressant drug-like actions in the vesicular monoamine transporter 2 (VMAT2) heterozygous (HET) mice (see Fukui et al., 2007). Note, these mutants were selected for study because they phenocopy the hypertensive patients that were treated with the VMAT inhibitor, reserpine, who experienced depression without anxiety (Freis, 1954), and because these findings were used as a basis for the monoamine hypothesis of depression (Schildkraut, 1965; Maes et al., 1994).

## 2 Results

### 2.1 Lisuride reduces motor activities in βArr1 and βArr2 mice

Locomotor, rearing, and stereotypical activities were examined at 5-min intervals across the 120 min test for the βArr1 (Supplementary Figures S1-3) and βArr2 mice (Supplementary Figures S4-6). For ease of viewing, the results are presented as cumulative motor activities at baseline (0-30 min) and following administration of the vehicle or various doses of lisuride (31-120 min). Overall, motor responses were reduced by lisuride in both genotypes.

When cumulative locomotor activities were examined at baseline for βArr1 animals, no significant genotype or treatment effects were found (Figure 1A). Following drug administration, overall cumulative locomotor activities were lower in βArr1-KO than in WT mice (*p*=0.046). With treatment, locomotion in both genotypes decreased in a dose-dependent fashion from 0.01 to 0.5 mg/kg lisuride relative to vehicle (*p*-values≤0.005); no further reductions occurred at higher doses.

**Figure 1.**
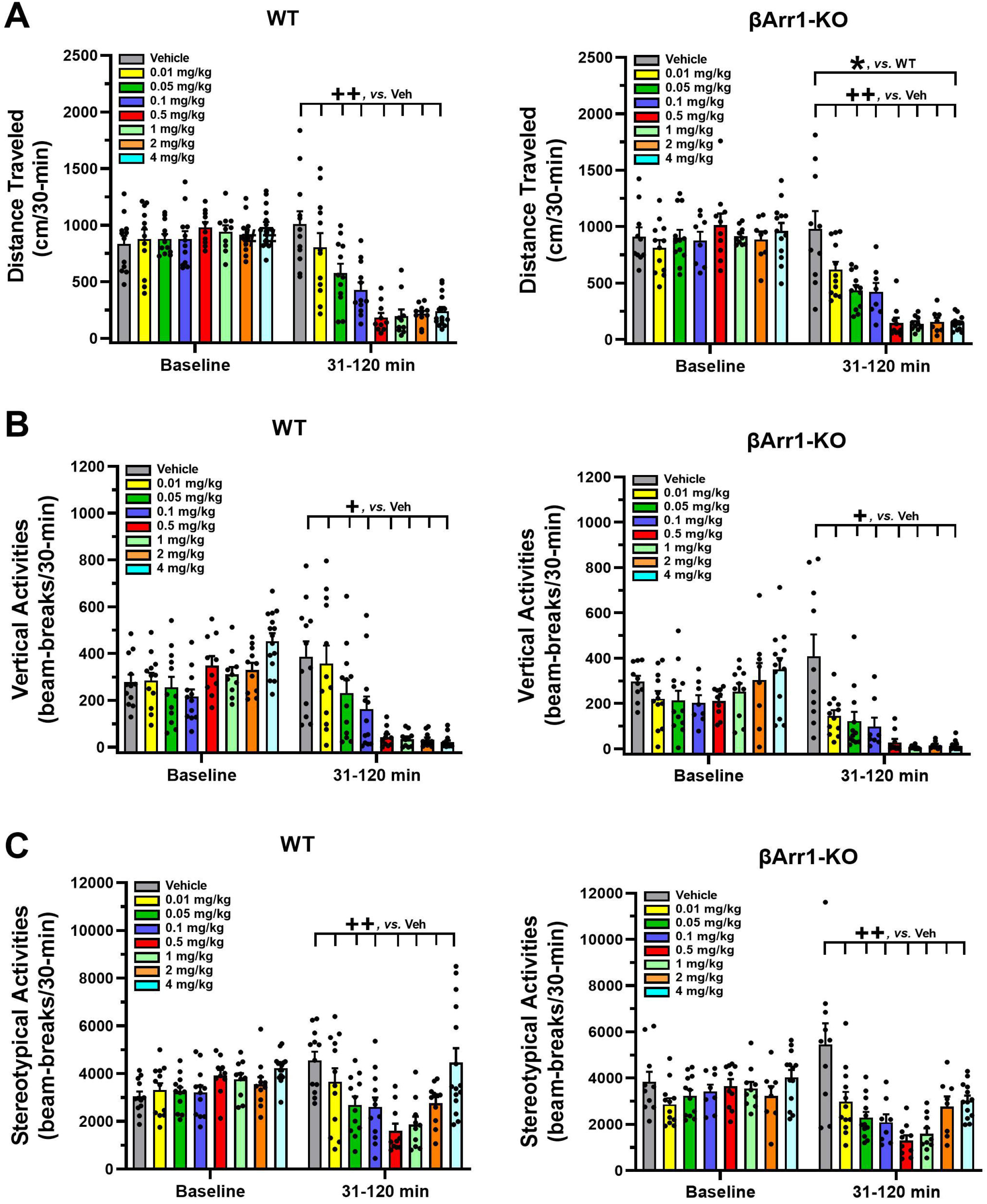
Effects of lisuride on cumulative motor activities in β-arrestin 1 mice. Baseline activities were monitored from 0-30 min, animals were given the vehicle (Veh) or various doses of lisuride, and returned to the open field for 90 min. **(A)** Cumulative distance traveled. Baseline: no significant effects. Post-injection: two-way ANOVA for genotype [F(1,161)=4.050, *p*=0.046] and treatment [F(7,161)=33.270, *p*<0.001]. **(B)** Vertical counts (rearing). Baseline: two-way ANOVA for genotype [F(1,161)=7.636, *p*=0.006] and treatment [F(7,161)=5.384, *p*<0.001]. Post-administration: two-way ANCOVA for treatment [F(7,160)=27.412, *p*<0.001]. **(C)** Stereotypical activities. Baseline: two-way ANOVA for treatment [F(7,161)=3.298, *p*=0.003]. Following injection: two-way ANCOVA for treatment [F(7,160)=14.609, *p*<0.001]. The data are presented as means ±SEMs. N=8-15 mice/genotype/treatment; \**p*<0.05, WT *vs.* KO; ^++^*p*<0.01, ^+^*p*<0.05, *vs.* Veh.

An examination of cumulative rearing activity found basal responses among genotypes (*p*=0.006) and the assigned treatment groups to be significantly different (*p*<0.001) (Figure 1B). To control for these conditions, post-administration cumulative rearing activities were subjected to analysis of covariance (ANCOVA). No significant genotype effects were detected; however, the initial suppression in rearing was visually more apparent in βArr1-KO than WT animals. Rearing activities in both genotypes decreased dose-dependently from vehicle to 0.05 mg/kg lisuride (*p*-values≤0.036). No additional reductions were present at higher doses.

Basal cumulative stereotypical activities were different also among groups prior to treatment (*p*=0.003) (Figure 1C). While no genotype effects at post-injection were noted with ANCOVA, cumulative stereotypies assumed a U-shaped function for dose with a decline from the vehicle through 0.01 to 0.5 mg/kg lisuride (*p*-values≤0.009) and then these activities increased with 4 mg/kg lisuride but were still below that of the vehicle control (*p*<0.008).

Motor activities were evaluated also in βArr2 mice. No genotype or treatment effects were discerned for baseline cumulative locomotor responses (Figure 2A). Nonetheless, post-administration a lisuride-induced depression in cumulative locomotor responses was more robust in βArr2-KO mice than in WT controls (*p*<0.001). All doses of lisuride suppressed locomotion compared to the vehicle (*p*-values<0.001) and they were not distinguished from each other.

**Figure 2.**
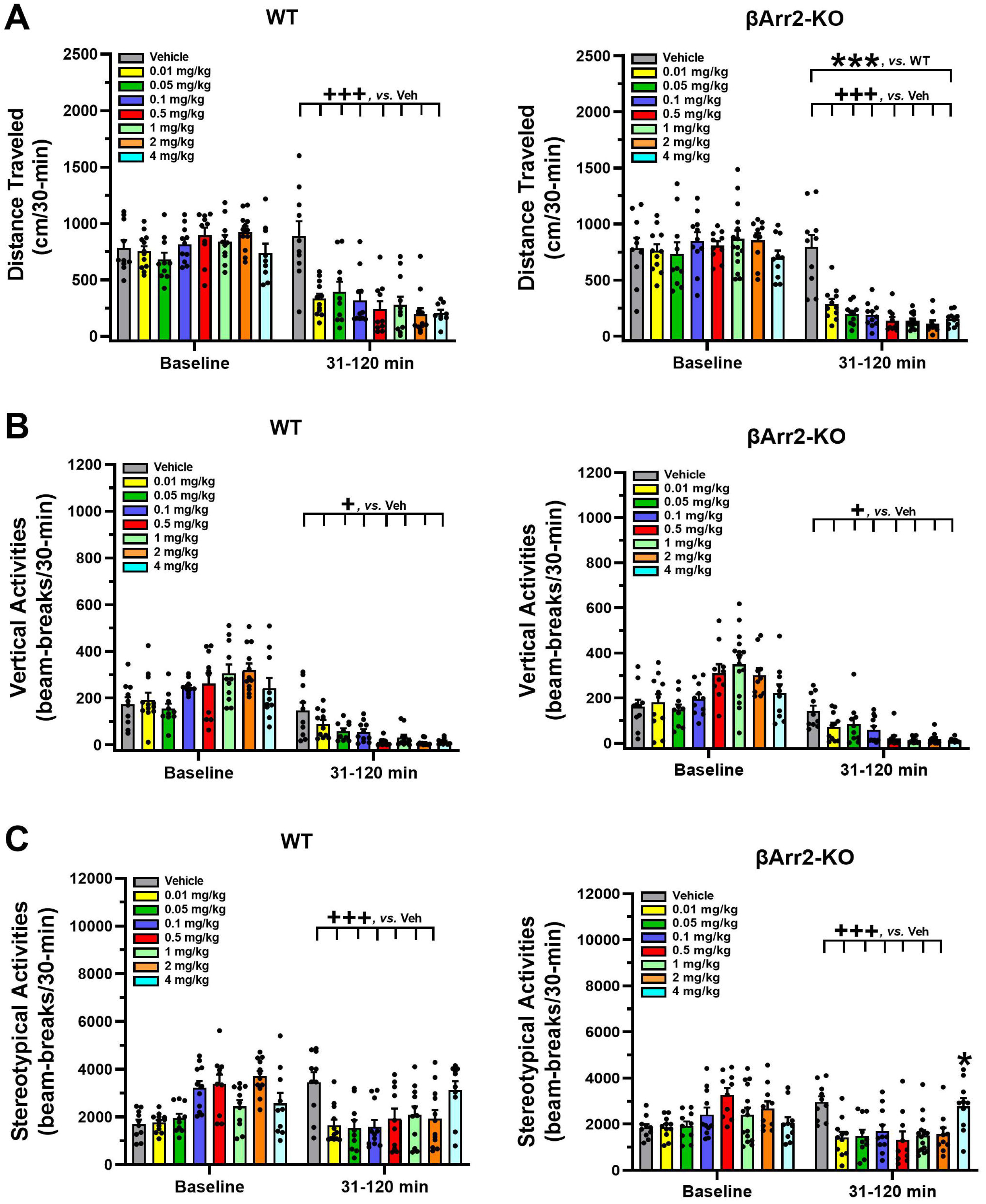
Effects of lisuride on cumulative motor activities in β-arrestin 2 mice. The procedure is identical to that described in Figure 1. **(A)** Cumulative distance traveled. Baseline: no significant effects. Post-injection: two-way ANOVA for genotype [F(1,157)=11.710, *p*<0.001] and treatment [F(7,157)=25.825, *p*<0.001]. **(B)** Vertical counts (rearing). Baseline: two-way ANOVA for treatment [F(7,157)=8.162, *p*<0.001]. Post-administration: two-way ANCOVA for treatment [F(7,156)=18.319, *p*<0.001]. **(C)** Stereotypical activities. Baseline: two-way ANOVA for genotype [F(1,157)=4.205, *p*=0.042] and treatment [F(7,157)=10.009, *p*<0.001]. Following injection: two-way ANCOVA for treatment [F(7,156)=8.485, *p*<0.001]. The data are presented as means ±SEMs. N=10-15 mice/genotype/treatment. \*\*\**p*<0.001, WT *vs.* KO; ^+++^*p*<0.001, ^+^*p*<0.05, *vs.* Veh.

An assessment of cumulative rearing activities in βArr2 animals revealed basal responses to be different across treatment assignments (*p*<0.001) (Figure 2B). The ANCOVA determined post-administration cumulative rearing to be depressed from the vehicle though all lisuride doses (*p*-values≤0.028)

Basal cumulative stereotypical activities were differentiated by genotype (*p*=0.042) and treatment assignment (*p*<0.001) (Figure 2C). Following injection of the vehicle or different doses of lisuride, ANCOVA discerned a trend for a genotype effect (*p*=0.056) where responses were lower overall in βArr2-KO than WT mice. Treatment effects were more dramatic and were U-shaped. Here, the numbers of cumulative stereotypical activities declined dramatically from the vehicle to 0.01 lisuride (*p*<0.001), the responses were flat to the 2 mg/kg dose (*p*-values≤0.001), then ascended with the 4 mg/kg dose to the level of the vehicle control.

Taken together, these findings show that in both βArr1 and βArr2 mice lisuride reduces cumulative locomotor, rearing, and stereotypical activities; however, these latter activities assume a biphasic U-shaped function with lisuride.

### 2.2 Lisuride effects on ethological behaviors in βArr1 and βArr2 mice

When HTRs were examined, both WT and βArr1-KO mice responded similarly to lisuride (Figure 3A). Nonetheless, the 4 mg/kg dose stimulated more HTRs than the vehicle and all other lisuride doses (*p*-values≤0.006). However, even with this high dose less than 7 HTRs were identified over the 30 min post-injection period. By comparison, in βArr2 animals no significant treatment effects were present where the numbers of HTRs barely exceeded 3 (Figure 3B). Thus, with lisuride HTRs are exceedingly low in both lines of mice.

**Figure 3.**
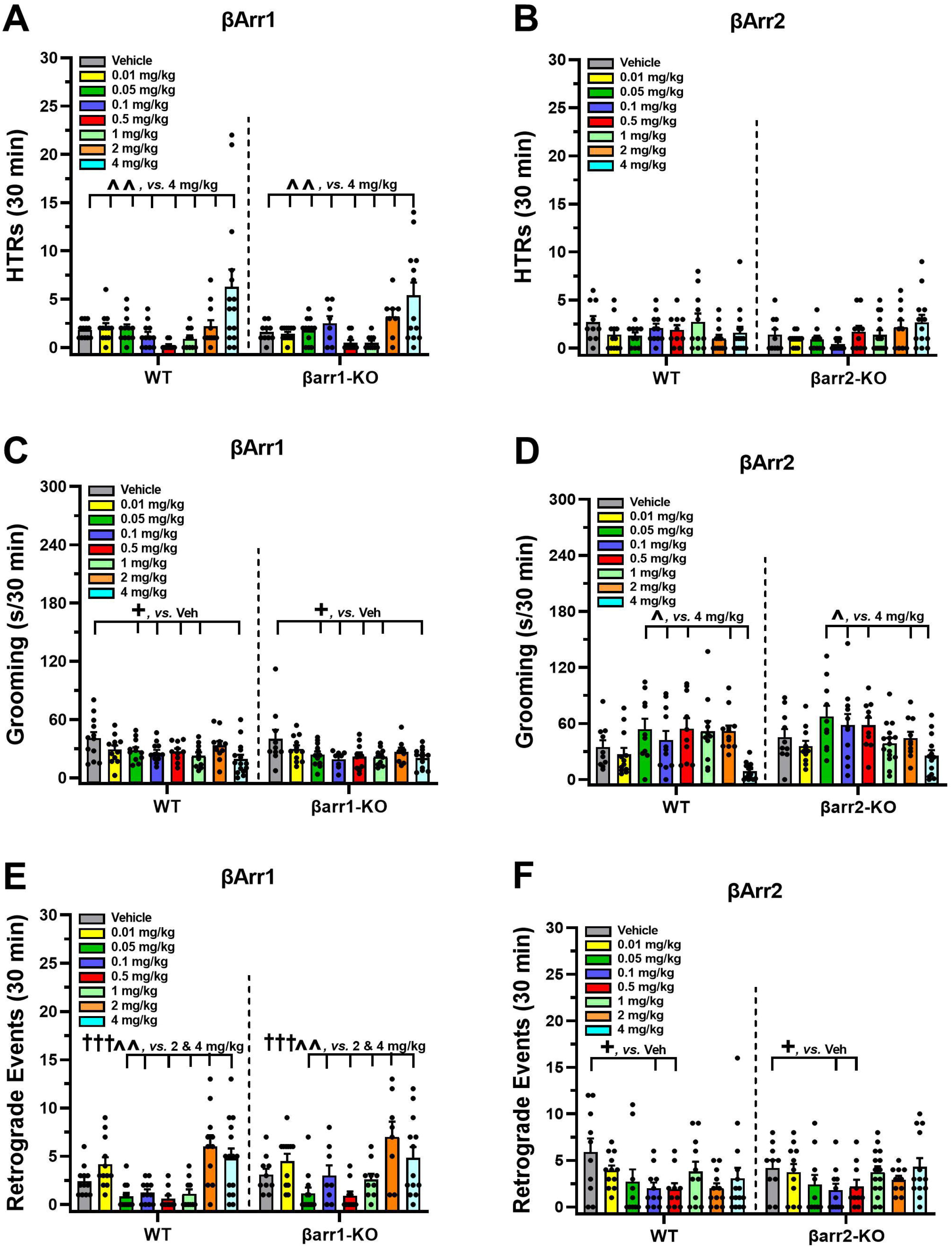
Effects of lisuride on ethological responses in β-arrestin 1 and β-arrestin 2 mice. Ethological behaviors were monitored in the open field over 30 min after injection of the vehicle or lisuride. **(A)** Head twitch responses in βArr1 mice. Two-way ANOVA for treatment [F(7,161)=9.519, *p*<0.001]. **(B)** Head twitch responses in βArr2 mice. Not significant. **(C)** Grooming in βArr1 mice. Two-way ANOVA for treatment [F(7,161)=4.438, *p*<0.001]. **(D)** Grooming in βArr2 mice. Two-way ANOVA for treatment [F(7,157)=4.206, *p*<0.001]. **(E)** Retrograde walking in βArr1 mice. Two-way ANOVA for treatment [F(7,161)=12.297, *p*<0.001]. **(F)** Retrograde walking in βArr2 mice. Two-way ANOVA for treatment [F(7,157)=3.385, *p*=0.002]. The data are presented as means ±SEMs. N=8-15 βArr1 mice/treatment; N=10-15 βArr2 mice/treatment. ^+^*p*<0.05, *vs.* Veh; ^^*p*<0.01, ^*p*<0.05, *vs.* 4 mg/kg lisuride; ^†††^*p*<0.001, *vs.* 2 mg/kg lisuride.

For self-grooming, no genotype effects were detected in either strain. Compared to the vehicle, the time spent grooming in βArr1 mice was decreased with 0.05 mg/kg lisuride and was maintained at a low level across doses to 1 mg/kg and then to the 4 mg/kg dose (*p*-vaules≤0.030) (Figure 3C). By dramatic contrast, in βArr2 animals lisuride produced an inverted U-shaped function for grooming. From a low at 0.01 mg/kg lisuride, grooming increased with the 0.05 mg/kg dose (*p*=0.021) and it remained high to 0.5 mg/kg then decreased with the highest dose (*p*-values≤0.048) (Figure 3D).

For retrograde walking, with lisuride the βArr1-KO tended to engage in more overall responses than WT controls (*p*=0.067) (Figure 3E). Retrograde walking with this drug assumed a biphasic U-shaped curve across doses. Lisuride decreased this behavior from the lowest dose to the 0.05 and 0.5 mg/kg doses (*p*-values<0.001), whereas 2 and 4 mg/kg lisuride augmented the response from the 0.5 to the 1 mg/kg doses (*p*-values≤0.021). When βArr2 mice were considered, a biphasic dose-response curve was found (Figure 3F). Significant declines from the vehicle were seen with the 0.1 and 0.5 mg/kg doses (*p*-values≤0.053), while retrograde walking events were augmented from this nadir with 4 mg/kg lisuride.

Collectively, only treatment effects with lisuride are obtained for ethological responses. Overall HTRs are higher in βArr1 than βArr2 mice, but they are still very low. The duration of grooming is decreased in βArr1 mice, but is enhanced in βArr2 animals except at the highest dose. By contrast, the incidences of retrograde walking are represented by a U-shaped function in both strains of mice.

### 2.3 Prepulse inhibition is reduced by lisuride in βArr1 but not in βArr2 mice

Effects of lisuride were analyzed for prepulse inhibition (PPI) in both βArr mouse strains. No genotype effects were detected with βArr1 mice (Figure 4A). Nonetheless, 0.5 mg/kg lisuride reduced PPI overall relative to vehicle and the 0.05 mg/kg group (*p*-values≤0.029). In contrast, no significant genotype or treatment effects were identified with βArr2 mice (Figure 4D).

**Figure 4.**
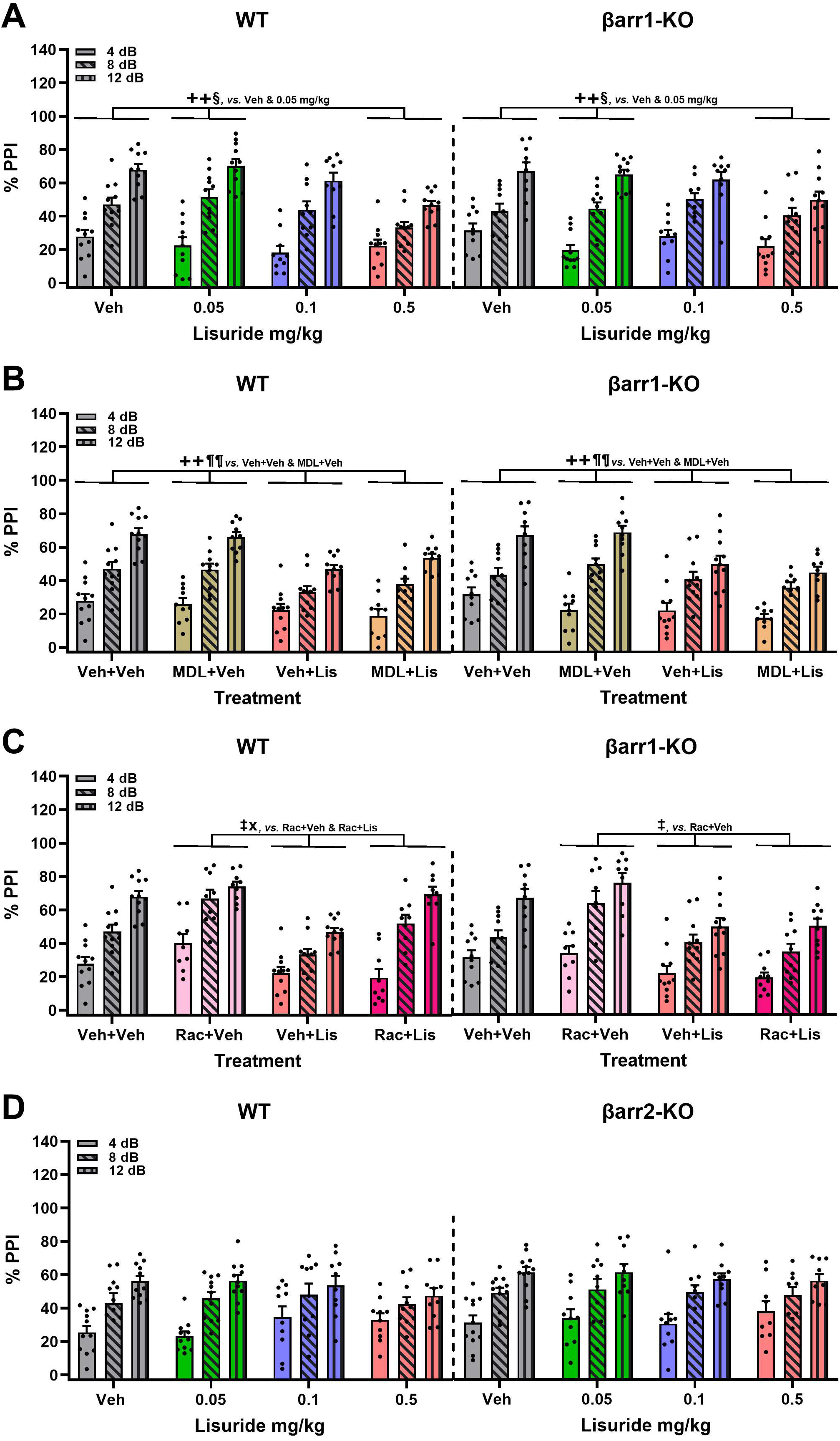
Effects of lisuride on prepulse inhibition in β-arrestin 1 and β-arrestin 2 mice. Mice were administered the MDL100907 or raclopride 15 min before giving the vehicle or lisuride and tested 25 min later. **(A)** PPI in βArr 1 mice receiving the vehicle or different doses of lisuride. RMANOVA for PPI [F(2,154)=369.848, *p*<0.001], PPI by treatment interaction [F(6,154)=5.964, *p*<0.001], and treatment [F(3,77)=4.579, *p*=0.005]. N=10-11 βArr1 mice/treatment. **(B)** PPI in βArr1 mice treated with the vehicle, MDL100907, or 0.5 mg/kg lisuride. RMANOVA for PPI [F(2,148)=327.131, *p*<0.001], PPI by treatment interaction [F(6,148)=4.611, *p*<0.001], and treatment [F(3,74)=9.722, *p*<0.001]. N=9-11 βArr1 mice/treatment. **(C)** PPI in βArr1 mice treated with the vehicle, raclopride, or 0.5 mg/kg lisuride. RMANOVA for PPI [F(2,144)=326.300, *p*<0.001], PPI by treatment interaction [F(6,144)=5.483, *p*<0.001], PPI by genotype by treatment interaction [F(6,144)=2.169, *p*=0.049], and treatment [F(3,72)=13.104, *p*<0.001]. N=9-11 βArr1 mice/treatment. **(D)** PPI in βArr 2 animals given the vehicle or different doses of lisuride. RMANOVA for PPI [F(2,150)=132.746, *p*<0.001] and PPI by treatment interaction [F(6,150)=2.485, *p*=0.025]. N=9-12 βArr2 mice/treatment. The data are presented as means ±SEMs. ^++^*p*<0.01, *vs.* Veh+Veh; ^§^*p*<0.05, *vs.*0.05 mg/kg lisuride; ^¶¶^*p*<0.01, *vs.* MDL+Veh; ^‡^*p*<0.05, *vs.* Rac+Veh; ^x^*p*<0.05, *vs.* Rac+Lis.

To determine whether the disruptive effect on PPI with lisuride was due to alterations in 5-HT2AR- or dopamine D2 receptor responses, βArr1 animals were treated with the respective antagonists, MDL100907 or raclopride. No genotype effected were noted. PPI with the vehicle and 0.5 mg/kg MDL100907 were virtually identical, while 0.5 mg/kg lisuride reduced PPI relative to both treatments (*p*-values≤0.007) (Figure 4B). When the 5-HT2AR antagonist and lisuride were given, the combined treatment failed to normalize PPI relative to the vehicle or MDL100907 groups (*p*-values≤0.002).

To determine if lisuride’s effects on PPI were mediated through D2 dopamine receptors, raclopride was tested next in βArr1 mice (Figure 4C). Compared to 3 mg/kg raclopride, PPI at 8 and 12 dB were decreased with 0.5 mg/kg lisuride in WT *versus* βArr1-KO mice (*p*-values≤0.035). Within WT animals, lisuride reduced PPI relative to raclopride at all prepulse intensities (*p*-values≤0.044) and at 12 dB with vehicle (*p*=0.003). Raclopride antagonism of lisuride’s effects normalized PPI. In βArr1-KO mice, effects on PPI were similar to that of WTs where PPI was reduced with lisuride at the 8 and 12 dB prepulses compared to raclopride (*p*-values≤0.006) and at 12 dB relative to vehicle (*p*=0.028). However, raclopride’s blockade of lisuride effects was similar to that of lisuride alone and this combined treatment was unable to restore PPI compared to raclopride at all three prepulses (*p*-values≤0.039) and with the vehicle at 12 dB (*p*=0.042). Besides PPI, null and startle activities were examined and only minor effects were detected within βArr1 lines and treatments (Supplementary Figure S7).

In summary, PPI is disrupted in βArr1 mice given 0.5 mg/kg lisuride and MDL100907 is unable to reverse this effect. Raclopride normalizes the lisuride-induced disruption of PPI, but only in WT mice; PPI in βArr1-KO animals is still abnormal.

### 2.4 Lisuride exerts anti-depressant drug-like actions in VMAT2 heterozygotes

WT and VMAT2-HET mice were used to assess possible anti-depressant drug-like actions of lisuride. Parenthetically Fukui and colleagues (2007) reported adult VMAT2-HETs were hypoactive in the open field, they presented with anhedonia-like responses to sucrose solutions, and showed enhanced immobility in tail suspension and forced swim that were normalized with tricyclic anti-depressants, selective serotonin and norepinephrine transporter inhibitors, and the atypical anti-depressant bupropion. These mutants were more responsive also to stress and displayed enhanced learned helplessness compared to their WT controls.

In preparation for the sucrose preference test, WT and VMAT2-HET mice were tested on day 1 with water-water (W-W) pairings and showed no preference for one bottle or side-preference over the other (Figure 5A). On day 2 mice were given the vehicle and presented with a 0.5% sucrose-water (S-W) pairing. The WT animals displayed a strong preference for the sucrose solution compared to mutants’ (*p*<0.001), whose preference was similar to W-W on day 1. Following acute administration of 0.5 mg/kg lisuride on day 3, the VMAT2-HETs’ sucrose preference exceeded that of WT mice (*p*=0.003) and it was significantly higher from their selections on days 1 (W-W) and 2 (S-W) (*p*-values<0.001). On day 4, the mutants’ sucrose preference was not significantly different from WT mice, but it had declined from day 3. However, by days 5 and 6 sucrose preference in the VMAT2-HET mice had fallen from WT levels (*p*≤0.010) and from their day 3 preference (*p*-values<0.001), such that their sucrose preference was similar to W-W on day 1. One reason why sucrose preference could vary across days in mutants could be due to differences in intake. When total fluid consumption was monitored, no genotype differences were found across days (Figure 5B). However, total intake significantly increased in all mice from days 1-3 compared to days 4-6 (*p*-values≤0.002). Collectively, VMAT2-HET mice fail to show a preference for sucrose until they are given lisuride. This preference persists for 2 days and declines thereafter, whereas sucrose preference is maintained in WT animals. Importantly, these differences in sucrose preference cannot be attributed to differential fluid consumption between the genotypes.

**Figure 5.**
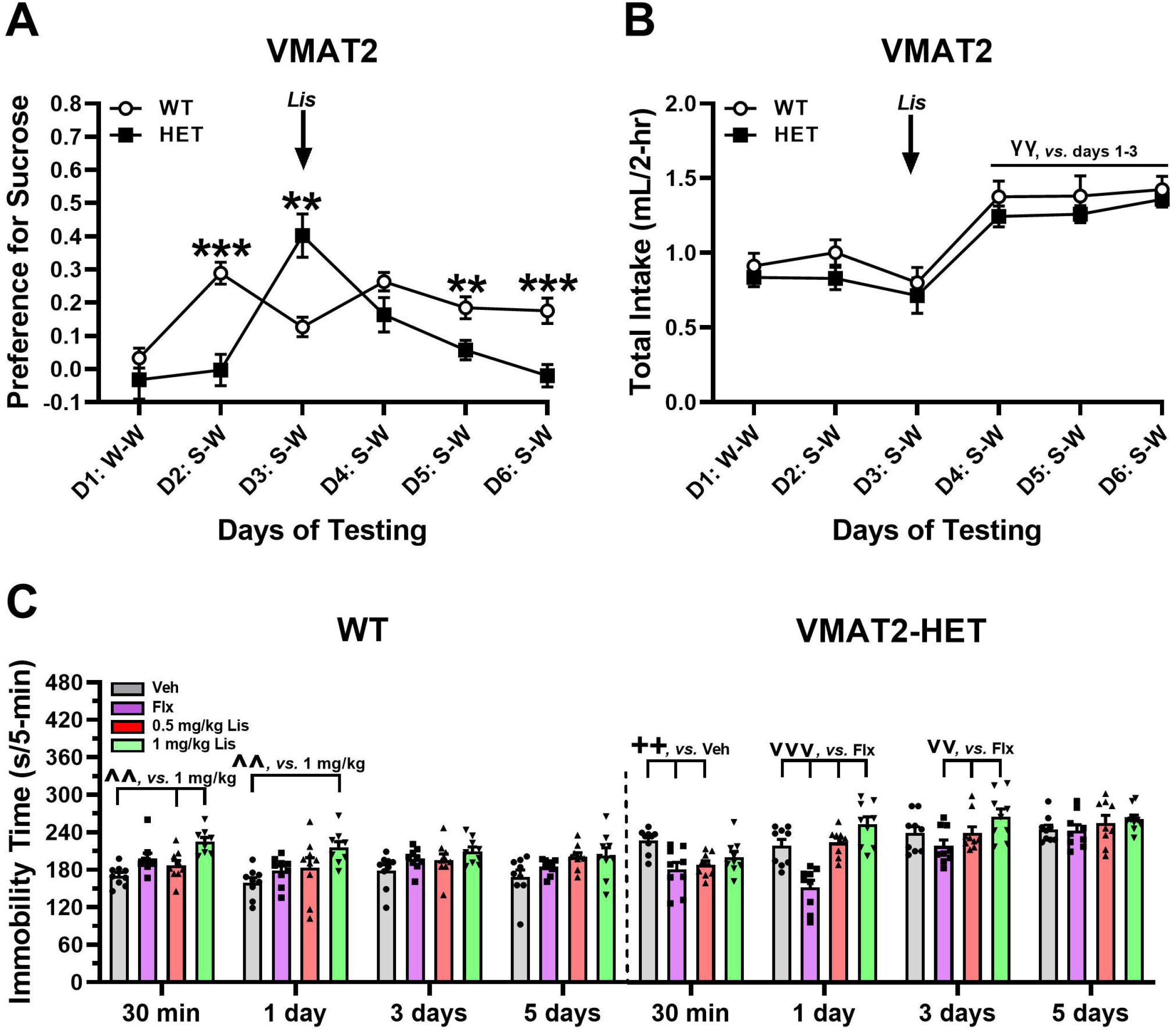
Effects of lisuride in tests of depressive-like behaviors in VMAT2 mice. In a two-bottle test, mice were presented with a water-water (W-W) pairing on day 1 and sucrose-water (S-W) pairings on days 2-6, and effects of the vehicle (Veh) and lisuride (Lis) were examined. **(A)** Sucrose preference in VMAT2 mice. RMANOVA for sucrose preference for day [F(5,110)=8.939, *p*<0.001], day by genotype interaction [F(5,110)=9.414, *p*<0.001], and genotype [F(1,22)=7.863, *p*=0.010]. **(B)** Total fluid intake. RMANOVA for the total liquid volume consumed for day [F(5,110)=21.585, *p*<0.001]. N=10-14 mice/genotype/treatment. **(C)** Tail suspension. RMANOVA for time in immobility by day [F(3,192)=17.424, *p*<0.001], day by genotype interaction [F(3,192)=17.452, *p*<0.001], day by treatment interaction [F(9,192)=3.538, *p*<0.001], day by genotype by treatment interaction [F(9,192)=2.783, *p*=0.004], genotype [F(1,64)=77.682, *p*<0.001], treatment [F(3,64)=14.256, *p*<0.001], and genotype by treatment [F(3,64)=7.557, *p*<0.001]. N=8-9 mice/genotype//treatment. All data are presented as means ±SEMs. \*\*\**p*<0.001, \*\**p*<0.01, WT *vs.* KO; ^γγ^*p*<0.01, *vs.* days 1-3; ^^*p*<0.01, *vs.* 1 mg/kg lisuride; ^++^*p*<0.01, *vs.* Veh; ^vvv^*p*<0.001. ^vv^*p*<0.01, *vs.* Flx.

In the tail suspension test, VMAT2 mice were treated acutely with the vehicle, 20 mg/kg fluoxetine, or 0.5 or 1 mg/kg lisuride and tested 30 min after injection and then over post-injection days 1, 3, and 5. Immobility in the VMAT2-HET mice treated with vehicle was significantly higher than in WT animals across all test days (*p*-values<0.001) (Figure 5C). Hence, repeated testing in the two genotypes was stable. In WT mice immobility times were not significantly different following acute treatment with fluoxetine or 0.05 mg/kg lisuride over all days of testing. Nevertheless, immobility times were increased acutely in WT animals with 1 mg/kg lisuride compared to vehicle and 0.5 mg/kg lisuride (*p*-values≤0.006), and these effects were maintained through post-injection day 1 relative to vehicle (*p*=0.002). In VMAT2-HETs, acute administration of fluoxetine or 0.5 mg/kg lisuride reduced immobility times compared to vehicle (*p*-values≤0.005). The effects of this lower lisuride dose were lost by day 1 post-injection, while those for fluoxetine persisted over that of the vehicle (*p*<0.001). By day 3 the fluoxetine effects were lost. In comparison, the 1 mg/kg lisuride was not efficacious on any of the test days in VMAT2-HET mice.

To determine if lisuride stimulated an increase in general motor activities that could confound the tail suspension results, VMAT2 mice were tested in the open field. In both WT and VMAT2-HET mice, 0.5 mg/kg lisuride depressed locomotor activities to similar extents in both genotypes relative to the vehicle control (Supplemental Figure S8). Thus, the antidepressant drug-like actions of lisuride cannot be attributed to an increase in general motor activity.

In summary, the tail suspension and sucrose preference results reveal that 0.5 mg/kg lisuride possesses anti-depressant drug-like actions that last acutely or at least 1 day later, respectively.

## 3. Discussion

In the present study, we examined responses to lisuride in WT, βArr1-KO, and βArr2-KO mice. Lisuride produced changes in many different responses in both βArr strains and in a few cases genotype effects were identified. For instance, overall locomotor activity was reduced by lisuride to a greater extent in mutants than WT animals. By comparison, no genotype effects in rearing and stereotypical activities were observed. The lisuride-induced HTRs were very low in all genotypes, even at the 4 mg/kg dose. With increasing dose, the time spent grooming was decreased in βArr1 mice while it assumed an inverted U-shaped function in βArr2 animals. Lisuride exerted no effect on PPI in βArr2 mice, whereas it disrupted PPI in both βArr1 genotypes. The 5-HT2AR antagonist MDL100907 failed to normalize PPI in the βArr1 strain, while the dopamine D2/D3 antagonist raclopride restored PPI in WT but not in βArr1-KO mice. Finally, lisuride was found to possess anti-depressant drug-like activities in VMAT2-HET mice, a mouse genetic model of depressive-like behaviors. Together, these results indicate that lisuride-induced responses were relatively uniform across the two stains of βArr mice with exceptions for grooming and PPI. In addition, while both the βArr1 and βArr2 strains were backcrossed extensively onto a C57BL/6J genetic background, the responses of WTs to lisuride in many tests were not fully concordant. The reason for this distinction between WTs is unclear at this time.

### 3.1 Motor performance

Locomotor, rearing, and stereotypical activities have been studied in various strains of rats--primarily males--and different strains of male and female mice. For locomotion, lisuride most commonly produced a biphasic response where low doses decreased and higher doses stimulated this activity (Carruba et al., 1980, 1985; Hara et al., 1982; Martin and Bendesky, 1984; Adams and Geyer, 1985; Fink and Morgenstern, 1985; Paulus and Geyer, 1991). Other researchers have observed decreases (Carruba et al., 1985; Nisoli et al., 1992; Chen et al., 2023) or increases in locomotion (Hlinak et al., 1983). Interestingly, when the biphasic effect in rats was followed over time in a behavioral pattern monitor (BPM), lisuride was reported to reduce locomotion over the first 30 min, but increase it over the final 30 min (Adams and Geyer, 1985). Biphasic responses have been reported also with mice in the open field with lisuride (Martin and Bendesky, 1984). By contrast, in mouse experiments in the BPM (Chen et al., 2023) or in our present studies locomotion was decreased in βArr mice with all lisuride doses. It should be emphasized that overall locomotion was decreased more in mutants than WTs from both βArr strains. Thus, it appears that βArr1 and βArr2 may play minor roles in the lisuride-induced suppression of locomotor activities in these mice.

In rats and mice, lisuride was reported to decrease rearing in the BPM and open field (Hlinak et al., 1983; Adams and Geyer, 1985; Chen et al., 2023). In studies with βArr mice, we found the same result. With respect to stereotypical activities, in rats lisuride was seen to elicit stereotyped sniffing (Hlinak et al., 1983; Carruba et al., 1985; Gerber et al., 1985), and licking or biting (Carruba et al., 1985). In the BPM with rats, the numbers of hole-pokes were enhanced and the numbers of pokes within the same hole were augmented with increased lisuride doses (Adams and Geyer, 1985). In mice, licking, chewing, and gnawing were stimulated by lisuride (Horowski and Wachtel, 1976), while hole-poking in the BPM was suppressed (Chen et al., 2023). Besides these responses, in rats lisuride produced behaviors associated with serotonin syndrome (Silbergeld and Hruska, 1979; Gerber et al., 1985; Marona-Lewicka et al., 2002). In the present studies some behaviors associated with this syndrome appeared in the βArr lines at the lowest lisuride dose. In regards to stereotypies, both βArr strains displayed a biphasic U-shaped function to the different doses of lisuride. Here, the lisuride-induced effects appeared to be more salient in βArr1 than βArr2 mice. While the individual stereotypical behaviors contributing to the U-shaped function are unclear, they probably represent the appearance and disappearance of many responses across time with the different the lisuride doses.

### 3.2 Ethological behaviors

With respect to ethological behaviors, psychedelics are reported to stimulate HTRs, grooming, and retrograde walking in rats and mice (Corne and Pickering, 1967; Kyzar et al., 2016; Rodriguiz et al., 2021). In rodents, the HTR has been taken as a proxy for hallucinogenic-like activities in humans (Corne and Pickering, 1967). This substitution of behaviors is strengthened as deletion of *Htr2a* in mice abrogates this response (González-Maeso et al., 2007; Keiser et al., 2009). Moreover, the potency of psychedelics in the mouse HTR are highly correlated with the dose to elicit hallucinations in humans (Glennon et al., 1984; Halberstadt et al., 2020). In humans, lisuride is reported to produce hallucinogenic activities in a single patient with prolactinoma given 1-2 mg/day (Turner et al., 1984), in another patient treated for migraine headache receiving 0.025 mg three times a day (Somerville and Herrmann, 1978), in five patients with Shy-Drager syndrome administered 0.5-5 mg/day (Lees and Bannister, 1981), and occasionally in some patients with Parkinson’s disease (Schachter et al., 1979; Parkes et al., 1981; LeWitt et al., 1982; Vaamonde et al., 1991). However, in normal subjects lisuride is devoid of psychedelic effects up to 600 µg (Herrmann et al., 1977). In rats and mice, lisuride does not produce HTRs (Gerber et al., 1985; Halberstadt and Geyer, 2013) and in the present investigations, the numbers of HTRs were undifferentiated by genotype and they were exceedingly low in the βArr1 and βArr2 even up to 4 mg/kg lisuride.

With grooming, lisuride have been reported to suppress this response in rats (Ferrari et al., 1992). In the present studies, no genotype effects were detected with the βArr1 or βArr2 mice. In βArr1 animals, lisuride decreased groom time. By contrast, in βArr2 mice a biphasic inverted U-shaped function emerged. Here, grooming increased from the lowest lisuride dose and declined slowly to the highest dose. Currently, the reason for differences between these βArr strains is obscure as they are both on a C57BL/6J genetic background.

Retrograde walking was also analyzed. In both βArr lines, lisuride produced a biphasic U-shaped dose-response curve. In βArr1 and βArr2 mice, lisuride reduced retrograde walking from the vehicle control, responding remained low across doses, and then increased with the 2 and 4 mg/kg doses. In summary, no genotype differences were found for HTRs and retrograde walking, whereas grooming was reduced with lisuride in βArr1 animals while adopting an inverted U-shaped function in βArr2 mice.

### 3.3 Prepulse inhibition

Psychedelics disrupt PPI in rats (Sipes and Geyer, 1994, Johansson et al., 1995; Ouagazzal et al., 2001; Halberstadt and Geyer, 2010; Páleníček et al., 2010), mice (Rodriguiz et al., 2021), and humans (Vollenweider et al., 2007; Schmid et al., 2015). PPI is abnormal also in individuals with schizophrenia (Braff et al., 2001) and LSD-induced states have been reported by some investigators to bear similarities to early acute phases of psychosis (Geyer and Vollenweider, 2008). In rats, lisuride was found to disrupt PPI (Halberstadt and Geyer, 2010). However, while the 5-HT2AR antagonist MDL 11,939 was unable to counteract lisuride’s effects, PPI was normalized with the dopamine D2/D3 antagonist raclopride (Halberstadt and Geyer, 2010). In the present studies, PPI was unaffected with lisuride in the βArr2 mice. By contrast, in βArr1 animals the highest dose of lisuride disrupted PPI. As with rats, the 5-HT2AR antagonist MDL100907 could not normalize PPI in either βArr1 genotype. However, raclopride was efficacious in restoring PPI only in WTs; βArr1-KOs were still deficient. This result suggests that restoration of PPI with lisuride requires βArr1-mediated signaling through some G protein coupled receptor other than the 5-HT2AR and the D2/D3 receptor.

### 3.4 Lisuride as an anti-depressant

There are now several small scale clinical studies indicating that psilocybin and LSD can alleviate depression and the effects are rapid and long-lasting (Bonson and Murphy, 1996; Carhart-Harris et al., 2016a, 2018, 2021; Griffiths et al., 2016; Ross et al., 2016; Reiff et al., 2020; Davis et al., 2021; Goodwin et al., 2022, 2023; McClure-Begley and Roth, 2022; von Rotz et al., 2022; Holze et al., 2023). Currently there are approximately 50 ongoing clinical trials with psilocybin, 2 trials with LSD, and additional trials with other psychedelics (see ClinicalTrials.gov). There is also intriguing evidence that lisuride may have anti-depressant actions in post-stroke patients experiencing depression (Hougaku et al., 1994). In C57BL/6J mice, acute restraint stress for 6 hr produced increased immobility in the forced swim and tail suspension tests and in both assays lisuride showed efficacy in decreasing these behaviors (Cao et al., 2022). It should be emphasized, these responses were tested only 30 min after drug administration and following a single 6 hr session of restraint stress. In the present investigation, we used a mouse genetic model of depressive-like behaviors to evaluate the effects of lisuride in the sucrose preference and tail suspension assays (see Fakui et al., 2007). In the S-W pairing, WT mice showed a strong preference for sucrose, whereas the VMAT2-HETs’ preference was not different from the W-W pairing. Hence, the mutants presented with anhedonia-like behavior. Following administration of lisuride, the preference for sucrose in the VMAT2-HET animals was robustly increased to a level higher than in WT controls. This preference in mutants was maintained through the next day, but it declined thereafter; the preference for sucrose was maintained in WTs across testing. Thus, in VMAT2-HET mice lisuride alleviated the anhedonia-like behavior at least for 2 days.

In tail suspension, VMAT2 mice were treated with the vehicle, fluoxetine, or 0.5 or 1 mg/kg lisuride and tested 30 min after injection, as well as, over post-injection days 1, 3, and 5. Immobility times in vehicle-treated VMAT2-HET animals were prolonged relative to WT controls and these differences were stable across days between genotypes. In WT mice, immobility times were stable with fluoxetine or 0.05 mg/kg lisuride, while the 1 mg/kg dose increased immobility over the first two days. By contrast, in VMAT2-HETs acute administration of fluoxetine or 0.5 mg/kg lisuride reduced immobility times *versus* the vehicle. These effects were transient for lisuride, while fluoxetine’s persisted through the next day. The 1 mg/kg dose was ineffective. The acute effect of lisuride on immobility time was specific and was not due to increased motor activity because locomotion in the open field was reduced dramatically with 0.5 mg/kg lisuride in the βArr1, βArr2, and VMAT2 mice. Together, lisuride relieved the behavioral despair acutely and the anhedonia-like behavior for at least for 2 days in the VMAT2-HET mice.

### 3.5 Biased signaling and behavior

Lisuride and LSD bind with high affinities to 5-HT2ARs (Hoyer et al., 1994) and both ligands are partial 5-HT2AR agonists (Egan et al., 1998; Kurrasch-Orbaugh et al., 2003; Zhang et al., 2022). Despite these similarities, LSD is hallucinogenic (Greiner et al., 1958), while lisuride is typically devoid of these effects in normal subjects (Herrmann et al., 1977). An interesting feature of LSD is that its psychedelic effects are long-lasting. This result may be attributed to the ability of the 5-HT2AR to “trap” LSD by forming a “lid” over the binding pocket containing LSD and, thereby, greatly slowing the dissociation rate of this ligand from the receptor (Wacker et al., 2017; Kim et al., 2020). Since lisuride is administered on a daily basis to parkinsonian patients (Schachter et al., 1979; Parkes et al., 1981; LeWitt et al., 1982; Vaamonde et al., 1991), it is unlikely this ligand is “trapped” in the 5-HT2AR like LSD--especially as its anti-depressant drug-like actions persist at most for 2 days in our studies.

Recently, the receptor structures of the 5-HT_2_ family have been reported with LSD, psilocin, and lisuride (Kim et al., 2020; Cao et al., 2022; Gumpper et al., 2022). Both ergolines and psilocin bind the receptors at the bottom of the orthosteric binding pocket in similar manners. However, these ligands bind within the extended binding pocket in subtly different ways that can produce slightly different receptor poses. For instance, the differential ergoline engagement with the Y370 residue in the extended binding pocket may affect transmembrane 7 positioning, thereby modulating βArr-mediated-signaling through the 5-HT2AR. Nonetheless, LSD activates G_q_, G_11_, and G_15_ proteins, with some activity at G_z_, and minimal activities at G_i_, G_12/13_, and G_s_ proteins (Kim et al., 2020). Similarly, lisuride stimulates G_q/11_ activation with calcium release (Cussac et al., 2008). Both βArr1 and βArr2 proteins can be recruited to the 5-HT2AR *in vitro* and, in cortical neurons, these βArrs are complexed to the receptor *in vivo* (Gelber, 1999). Recently, it has been demonstrated that LSD is βArr biased at the 5-HT2AR *in vitro* (Wacker et al., 2013; Wang et al., 2013; Kim et al., 2020) and βArr2 biased *in vivo* (Rodriguiz et al., 2021). By comparison, results from the present investigations in βArr1-KO and βArr2-KO mice indicate that lisuride is G protein biased at the 5-HT2AR since deletion of *Arrb1* or *Arrb2* results in very low incidences of HTRs--even with the 4 mg/kg dose. By comparison, LSD robustly stimulates HTRs in βArr1-KO and WT animals from both βArr lines of mice; this effect is lost in βArr2-KOs (Rodriguiz et al., 2021).

Since LSD is βArr2 biased while promoting some G protein-mediated activities at the 5-HT2AR (Wacker et al., 2017; Kim et al., 2020) and because LSD is reported to exert anti-depressant actions (Gasser et al., 2015), we tested whether the G-protein biased ligand lisuride would have anti-depressant drug-like actions and confirmed this response in the VMAT2-HET mice. A possible limitation of these experiments is that lisuride binds many dopaminergic, adrenergic, and other serotonergic receptors (Piercey et al., 1996; Egan et al., 1998; Marona-Lewicka et al., 2002; Millan et al., 2002; Kroeze et al., 2015). Hence, its effects could be attributed to actions at other receptors or at their combinations of actions. A recent publication addresses this conundrum where a novel G protein biased 5-HT2AR compound was tested in the VMAT2 and in learned helplessness models with the sucrose preference and/or tail suspension tests (Kaplan et al., 2022). The biased compound stimulated very low HTRs and was not reinforcing in conditioned place preference. Importantly, in VMAT2-HETs it showed anti-depressant drug-like actions in tail suspension that lasted for 2 days--the maximum tested. By comparison, in learned helplessness a single administration of the G protein biased compound lasted at least 3 days in sucrose preference and this was similar to that for psilocin. Immobility in tail suspension was reduced with the compound for 14 days, while psilocin was efficacious for 9 days. The results from these studies and the present investigation with lisuride suggest that G protein biased signaling at the 5-HT2AR may represent a novel way to separate the hallucinogenic-like potential of psychedelics from their anti-depressant drug-like actions.

## 4 Materials and methods

### 4.1 Animals

Adult male and female WT and global βArr1-KO (Kim et al., 2018), WT and global βArr2-KO (Bohn et al., 1999), and WT and VMAT2-HET mice (Wang et al., 1997) served as the subjects. All strains of mice had been backcrossed onto a C57BL/6J genetic background for more than 10 generations. All animals were housed 3-5 per cage according to sex and genotype on a 12:12 hr light:dark cycle (lights on 0700 hr) in a humidity- and temperature-controlled room, with food and water provided *ad libitum*. The behavioral experiments were conducted between 0900-1700 hr when the mice were 3-8 months of age. All experiments were conducted with a protocol approved by the Institutional Animal Care and Use Committee at Duke University and according to ARRIVE guidelines.

### 4.2 Drugs

The drugs consisted of lisuride (Sigma-Aldrich, St Louis, MO), fluoxetine (Sigma-Aldrich), MDL 100907, and raclopride (Biotechne Corporation, Minneapolis, MN). The vehicle consisted of *N,N*-dimethyllacetamide (final volume 0.5%; Sigma-Aldrich) and it was brought to volume with 5% 2-hydroxypropoyl-β-cyclodextrin (Sigma-Aldrich) in water (Mediatech Inc., Manassas, VA). All drugs were injected (i.p.) in a 5 ml/kg volume.

### 4.3 Open field activity

The open field consisted of a clear Plexiglas arena 21 × 21 × 30 cm (Omnitech Electronics, Columbus, OH) that was illuminated at 180 lux (Rodriguiz et al., 2021). Overhead cameras recorded behaviors. Mice were placed into the open field, 30 min later they were injected with the vehicle or different doses of lisuride, and immediately returned to the open field for 90 min. Fusion Integra software (Omnitech) recorded locomotor activity (distance traveled), rearing (vertical beam-breaks), and stereotypical activities (repetitive beam-breaks less than 1 sec) in 5-min blocks.

### 4.4 Head twitch, grooming, and retrograde walking

These behaviors were filmed during the test for motor activity (Rodriguiz et al., 2021). All responses were scored over the first 30 min following administration of the vehicle or lisuride after collection of baseline activity. The HTRs were scored by observers blinded to the sex, genotype, and treatments of the mice. Grooming and retrograde walking were scored by the TopScan program (Clever Sys Incorporated, Reston, VA). The data are presented as the number of head twitches, time spent grooming, and the numbers of retrograde walking events.

### 4.5 Prepulse inhibition (PPI)

PPI of the acoustic startle response was monitored in SR-LAB chambers (San Diego Instruments, San Diego, CA) (Rodriguiz et al., 2021). Mice were administered the vehicle or different doses of lisuride and returned to their home cages. When the vehicle, MDL100907, or raclopride were given, they were administered 15 min prior to subsequent injection with the vehicle or lisuride. Following the vehicle or lisuride injection, the animals were placed into the apparatus. After 10 min of habituation to a white noise background (64 dB) a series of 42 trials was given. Trials consisted of either 6 null trials, 18 pulse-alone trials, or 18 prepulse-pulse trials. Null trials consisted of the white noise background, pulse trials were composed of 40 msec bursts of 120 dB white-noise, and prepulse-pulse trials consisted of 20 msec pre-pulse stimuli that were 4, 8, or 12 dB above the white-noise background (6 trials/dB), followed 80 msec later by the 120 dB pulse stimulus. Testing commenced with 10 pulse-alone trials followed by combinations of the prepulse-pulse and null trials, and was terminated with 10 pulse-alone trials. PPI responses were calculated as %PPI = [1–(pre-pulse trial Amplitude/startle-only trial Amplitude)]*100.

### 4.6 Sucrose preference

VMAT2 mice were housed individually for 7 days prior to and throughout the study (Fukui et al., 2007; Kaplan et al., 2023). Food was available *ad libitum*, while the water bottle was removed 2.5 hr from the home-cage before the beginning of the dark cycle. A pair of bottles was filled with water and the procedure was repeated until water consumption was stable and no side-bias was evident. Subsequently, each pair of bottles was filled with water or 0.5% sucrose. Note, the solution was prepared just prior to testing, the bottles were weighed before and immediately after the test, and they were cleaned and drained thoroughly each day. The pair of bottles was presented to the mice 1.5 hr into their dark cycle (4 hr of total fluid deprivation) and the animals were allowed to drink for 2 hr. After this time, the bottles were removed and the original water bottle was returned to the home-cage at least 1 hr later. The final water-water pairing occurred on day 1. The next day, animals were presented with the sucrose-water pairing. On day 3 they were administered an acute injection of 1 mg/kg lisuride (i.p.) and 5 min later were given the sucrose-water pairing. Subsequent sucrose-water pairings were presented over days 4-6. The total volume of liquid consumed in the 2 hr test was determined each day. Preference for the sucrose solution was calculated by dividing the volume of sucrose consumed by the total volume of liquid consumption.

### 4.7 Tail Suspension

This test was conducted in the morning. VMAT2 mice were administered the vehicle, fluoxetine, or lisuride and were tested 30 min later (see Fukui et al., 2007). Mice were suspended by the tail for 6 min using medical tape attached to the Med Associates (St. Albans, VT) tail suspension hook connected to a load cell on the ceiling of the chamber. The signal from the load cell was amplified and processed with Med-Associates Tail Suspension Software that determined immobility times. All load cells were calibrated prior to the start of testing each day according to the manufacturer’s instructions.

### 4.8 Statistics

The results are presented as means ±SEMs; all statistics were performed with SPSS 27 programs (IBM-SPSS, Chicago, IL). Univariate ANOVAs analyzed treatment effects. If cumulative baseline motor activities were significantly different for genotypes or for treatment assignments, the cumulative post-administration results were analyzed by analysis of covariance. Repeated measures ANOVA were used for analysis of open field, PPI, sucrose preference, and tail suspension studies. Bonferroni-corrected pair-wise comparisons were used as the *post-hoc* tests. A *p*<0.05 was considered statistically significant. The results displayed in all figures were plotted using GraphPad Prism 9.5.1 (GraphPad Software, Boston, MA).

## Data available statement

Data that support this study are available from the corresponding authors upon reasonable request.

## Ethics statement

All experiments were conducted with a protocol approved by the Institutional Animal Care and Use Committee at Duke University and according to ARRIVE guidelines.

## Author contributions

V.M.P. conducted the behavioral experiments and graphed the data with R.M.R and W.C.W. V.P.M. and R.M.R. statistically analyzed the data. W.C.W. and B.L.R. conceived the experiments, proposed the experimental designs, and wrote the manuscript.

## Acknowledgements

We wish to thank Dr. Robert J. Lefkowitz (Duke University Medical Center, Durham, NC, USA) for providing us with the new strain of βArr1 mice and Dr. Laura M. Bohn (Scripps Research Institute, Jupiter, FL, USA) for sending us the βArr2 mice. We thank Mr. Mitchell Huffstickler, Mr. Christopher Means, Ms. Julia Hoyt, Ms. Sarah Page Steffens, Ms. Ann Njoroge, and Mr. Nathan Franklin for scoring the videos for HTRs and helping with some of the behavioral experiments. We thank also Ms. Jiechun Zhou for breeding, genotyping, and maintaining the βArr1, βArr2, and VMAT2 mice. Some of the behavioral experiments were conducted with equipment and software purchased with a North Carolina Biotechnology Center grant. The work was supported by NIDA grant R37-DA045657 and DARPA [Grant DARPA-5822 (HR00112020029)]. The views, opinions, and/or findings contained in this material are those of the authors and should not be interpreted as representing the official views, policies, or endorsement of the Department of Defense or the U.S. Government.

