## Supplementary Figures and Legends for "The G protein biased serotonin 5-HT 2A receptor agonist lisuride exerts anti-depressant drug-like activities in mice"

**Supplementary materials**

*Supplementary figure legends*

**Figure S1**

Effects of lisuride in β-arrestin 1 mice on locomotor activities in 5-min segments at baseline (0-30 min) and after vehicle or lisuride injection (31-120 min). βArr1 mice were placed into the open field and 30 min later were given the vehicle or various doses of lisuride and returned immediately to the open field for 90 min. RMANOVA for baseline (0-30 min) for time [F(5,805)=165.478, *p*<0.001], time by genotype interaction [F(5,805)=2.335, *p*=0.041], and time by treatment interaction [F(35,805)=2.375, *p*<0.001]. RMANOVA for post-administration (31-120 min) for time [F(17,2737)=53.956, *p*<0.001], time by treatment interaction [F(119,2737)=3.589, *p*<0.001], genotype [F(1,161)=4.050, *p*=0.046], and treatment [F(7,161)=33.270, *p*<0.001]. The same vehicle results are presented in panels A-C and doses of lisuride are dispersed across panels for easier visualization of effects over time. **(A)** Locomotor activities in β-arrestin 1 animals administered the vehicle or 0.01, 0.5, or 4 mg/kg lisuride. **(B)** Locomotor responses in β-arrestin 1 mice given the vehicle or 0.05, or 2 mg/kg lisuride. **(C)** Locomotion in β-arrestin 1 subjects receiving the vehicle or 0.1, 1 mg/kg lisuride. The data are presented as means ±SEMs. N=8-15 mice/genotype/treatment.

**Figure S2**

Effects of lisuride in β-arrestin 1 mice on rearing activities in 5-min segments at baseline (0-30 min) and after vehicle or lisuride injection (31-120 min). The procedure and justification for presentation of groups in each panel is described in the legend for Figure S1. RMANOVA for baseline (0-30 min) for time [F(5,805)=37.360, *p*<0.001], time by genotype interaction [F(5,805)=7.630, *p*<0.001], time by treatment interaction [F(35,805)=3.034, *p*<0.001], genotype [F(1,161)=7.636, *p*=0.006], and treatment [F(7,161)=5.384, *p*<0.001]. RMANOVA for post-administration (31-120 min) for time [F(17,2737)=4.654, *p*<0.001], time by treatment interaction

[F(119,2737)=2.969, $p$<0.001], genotype [F(1,161)=5.746, $p$=0.018], and treatment [F(7,161)=19.819, $p$<0.001]. **(A)** Rearing activities in β-arrestin 1 animals administered the vehicle or 0.01, 0.5, or 4 mg/kg lisuride. **(B)** Rearing responses in β-arrestin 1 mice given the vehicle or 0.05, or 2 mg/kg lisuride. **(C)** Rearing in β-arrestin 1 subjects receiving the vehicle or 0.1, 1 mg/kg lisuride. The data are presented as means ±SEMs. N=8-15 mice/genotype/treatment.

**Figure S3**

Effects of lisuride in β-arrestin 1 mice on stereotypical activities in 5-min segments at baseline (0-30 min) and after vehicle or lisuride injection (31-120 min). The procedure and justification for presentation of groups in each panel is described in the legend for Figure S1. RMANOVA for baseline (0-30 min) for time [F(5,805)=114.282, $p$<0.001], time by genotype interaction [F(5,805)=2.196, $p$=0.053], time by treatment interaction [F(35,805)=1.829, $p$=0.003], and treatment [F(7,161)=3.298, $p$=0.003]. RMANOVA for post-administration (31-120 min) for time [F(17,2737)=40.066, $p$<0.001], time by treatment interaction [F(119,2737)=9.823, $p$<0.001], and treatment [F(7,161)=13.598, $p$<0.001]. **(A)** Stereotypical activities in β-arrestin 1 animals administered the vehicle or 0.01, 0.5, or 4 mg/kg lisuride. **(B)** Stereotypical responses in β-arrestin 1 mice given the vehicle or 0.05, or 2 mg/kg lisuride. **(C)** Stereotypies in β-arrestin 1 subjects receiving the vehicle or 0.1, 1 mg/kg lisuride. The data are presented as means ±SEMs. N=8-15 mice/genotype/treatment.

**Figure S4**

Effects of lisuride in β-arrestin 2 mice on locomotor activities in 5-min segments at baseline (0-30 min) and after vehicle or lisuride injection (31-120 min). The procedure and justification for presentation of groups in each panel is described in the legend for Figure S1. RMANOVA for baseline (0-30 min) for time [F(5,785)=185,310, $p$<0.001] and time by treatment interaction

[F(35,785)=1.909, *p*<0.001]. RMANOVA for post-administration (31-120 min) for time [F(17,2669)=37.609, *p*<0.001], time by treatment interaction [F(119,2669)=1.745, *p*<0.001], genotype [F(1,157)=11.710, *p*<0.001], and treatment [F(7,157)=25.825, *p*<0.001]. **(A)** Locomotor activities in β-arrestin 2 animals administered the vehicle or 0.01, 0.5, or 4 mg/kg lisuride. **(B)** Locomotor responses in β-arrestin 2 mice given the vehicle or 0.05, or 2 mg/kg lisuride. **(C)** Locomotion in β-arrestin 2 subjects receiving the vehicle or 0.1, 1 mg/kg lisuride. The data are presented as means ±SEMs. N=10-15 mice/genotype/treatment.

**Figure S5**

Effects of lisuride in β-arrestin 2 mice on rearing activities in 5-min segments at baseline (0-30 min) and after vehicle or lisuride injection (31-120 min). The procedure and justification for presentation of groups in each panel is described in the legend for Figure S1. RMANOVA for baseline (0-30 min) for time [F(5,785)=65.738, *p*<0.001], time by treatment interaction [F(35,785)=3.486, *p*<0.001], and treatment [F(7,157)=8.162, *p*<0.001]. RMANOVA for post-administration (31-120 min) for time [F(17,2669)=10.316, *p*<0.001], time by treatment interaction [F(119,2669)=2.279, *p*<0.001], and treatment [F(7,157)=16.459, *p*<0.001]. **(A)** Rearing activities in β-arrestin 2 animals administered the vehicle or 0.01, 0.5, or 4 mg/kg lisuride. **(B)** Rearing responses in β-arrestin 2 mice given the vehicle or 0.05, or 2 mg/kg lisuride. **(C)** Rearing in β-arrestin 2 subjects receiving the vehicle or 0.1, 1 mg/kg lisuride. The data are presented as means ±SEMs. N=10-15 mice/genotype/treatment.

**Figure S6**

Effects of lisuride in β-arrestin 2 mice on stereotypical activities in 5-min segments at baseline (0-30 min) and after vehicle or lisuride injection (31-120 min). The procedure and justification for presentation of groups in each panel is described in the legend for Figure S1. RMANOVA for baseline (0-30 min) for time [F(5,785)=107.945, *p*<0.001], time by treatment interaction

[F(35,785)=3.155, *p*<0.001], genotype [F(1,157)=4.205, *p*=0.042], and treatment [F(7,157)=10.009, *p*<0.001]. RMANOVA for post-administration (31-120 min) for time [F(17,2669)=29.380, *p*<0.001], time by treatment interaction [F(119,2669)=6.127, *p*<0.001], genotype [1,157]=4.031, *p*=0.046], and treatment [F(7,157)=8.744, *p*<0.001]. **(A)** Stereotypical activities in β-arrestin 2 animals administered the vehicle or 0.01, 0.5, or 4 mg/kg lisuride. **(B)** Stereotypical responses for β-arrestin 2 mice given the vehicle or 0.05, or 2 mg/kg lisuride. **(C)** Stereotypies in β-arrestin 2 subjects receiving the vehicle or 0.1, 1 mg/kg lisuride. The data are presented as means ±SEMs. N=10-15 mice/genotype/treatment.

**Figure S7**

Effects of lisuride on null and startle activities for PPI in β-arrestin 1 and β-arrestin 2 mice. See the legend in Figure 4 for details. **(A)** Null activities in βArr1 mice treated with vehicle or lisuride. Not significant. **(B)** Startle activities in βArr1 subjects with the same treatments as panel A. Two-way ANOVA for genotype [F(1,77)=9.511, *p*=0.003], treatment [F(3,77)=9.896, *p*<0.001], and genotype by treatment interaction [F(3,77)=2.952, *p*=0.038]. N=10-11 βArr1 mice/treatment. N=10-11 mice/genotype/treatment. **(C)** Null activities in βArr1 mice administered the vehicle, MDL100907, or lisuride. Two-way ANOVA for treatment [F(3,74)=3.294, *p*=0.025]. **(D)** Startle activities in βArr1 mice given the same treatments as panel C. Two-way ANOVA for treatment [F(3,74)=13.421, *p*<0.001]. N=9-11 βArr1 mice/treatment. **(E)** Null activities in βArr1 mice receiving the vehicle, raclopride, or lisuride. Two-way ANOVA for genotype [F(1,72)=4.213, *p*=0.044]. **(F)** Startle activities in βArr1 mice receiving identical treatments as panel E. Two-way ANOVA for genotype [F(1,72)=6.059, *p*=0.016], treatment [F(3,72)=9.301, *p*<0.001], and genotype by treatment interaction [F(3,72)=4.173, *p*=0.009]. N=9-11 βArr1 mice/treatment. **(G)** Null activities in βArr2 animals given vehicle or lisuride. Two-way ANOVA for genotype by treatment interaction [F(3,75)=2.781, *p*=0.047]. **(H)** Startle activities in βArr2 mice undergoing the same treatments as panel C. Two-way ANOVA for treatment [F(3,75)=7.356, *p*<0.001]. N=9-12 βArr2 mice/treatment. The data are presented as means ±SEMs. ***$p<0.001$, WT *vs.* KO; $^{++}p<0.01$, $^{+}p<0.01$, *vs.* Veh; $^{\wedge\wedge\wedge}p<0.001$, $^{\wedge\wedge}p<0.01$, $^{\wedge}p<0.05$, *vs.* 0.5 mg/kg lisuride or Veh+Lis; $^{\P}p<0.05$, *vs.* MDL+Veh; $^{x}p<0.05$, *vs.* Rac+Lis.

**Figure S8**

Effects of lisuride in VMAT2 mice on locomotor activities in 5-min segments at baseline (0-30 min) and after vehicle or lisuride injection (31-120 min). The procedure is described in the legend for Figure S1. **(A)** RMANOVA for baseline (0-30 min) for time [$F(5,100)=15.555$, $p<0.001$]. RMANOVA for post-administration (31-120 min) for time [$F(17,340)=10.327$, $p<0.001$], time by treatment interaction [$F(17,340)=3.631$, $p=0.002$], and treatment [$F(1,20)=99.297$, $p<0.001$]. **(B)** Effects of lisuride on cumulative motor activities in VMAT2 mice. Baseline activities were monitored from 0-30 min, animals were given the vehicle (Veh) or various doses of lisuride, and returned to the open field for 90 min. Cumulative distance traveled. Baseline: no significant effects. Post-injection: two-way ANOVA for treatment [$F(1,20)=99.297$, $p<0.001$]. The data are presented as means ±SEMs. N=6 mice/genotype/treatment. $^{+}p<0.05$, *vs.* Veh.

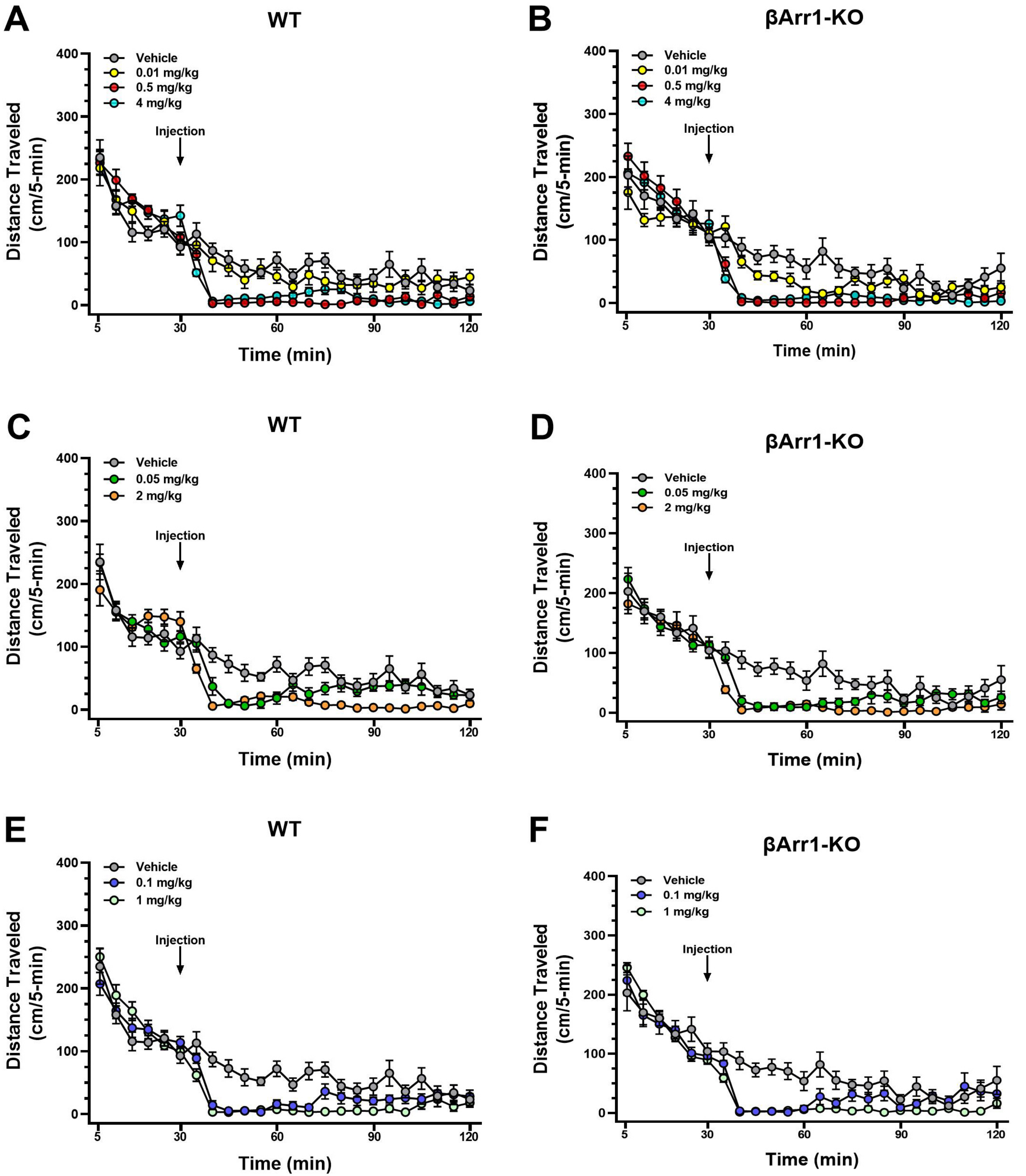

A
WT
Vehicle
0.01 mg/kg
0.5 mg/kg
4 mg/kg
Injection
Distance Traveled (cm/5-min)
Time (min)
B
βArr1-KO
Vehicle
0.01 mg/kg
0.5 mg/kg
4 mg/kg
Injection
Distance Traveled (cm/5-min)
Time (min)
C
WT
Vehicle
0.05 mg/kg
2 mg/kg
Injection
Distance Traveled (cm/5-min)
Time (min)
D
βArr1-KO
Vehicle
0.05 mg/kg
2 mg/kg
Injection
Distance Traveled (cm/5-min)
Time (min)
E
WT
Vehicle
0.1 mg/kg
1 mg/kg
Injection
Distance Traveled (cm/5-min)
Time (min)
F
βArr1-KO
Vehicle
0.1 mg/kg
1 mg/kg
Injection
Distance Traveled (cm/5-min)
Time (min)

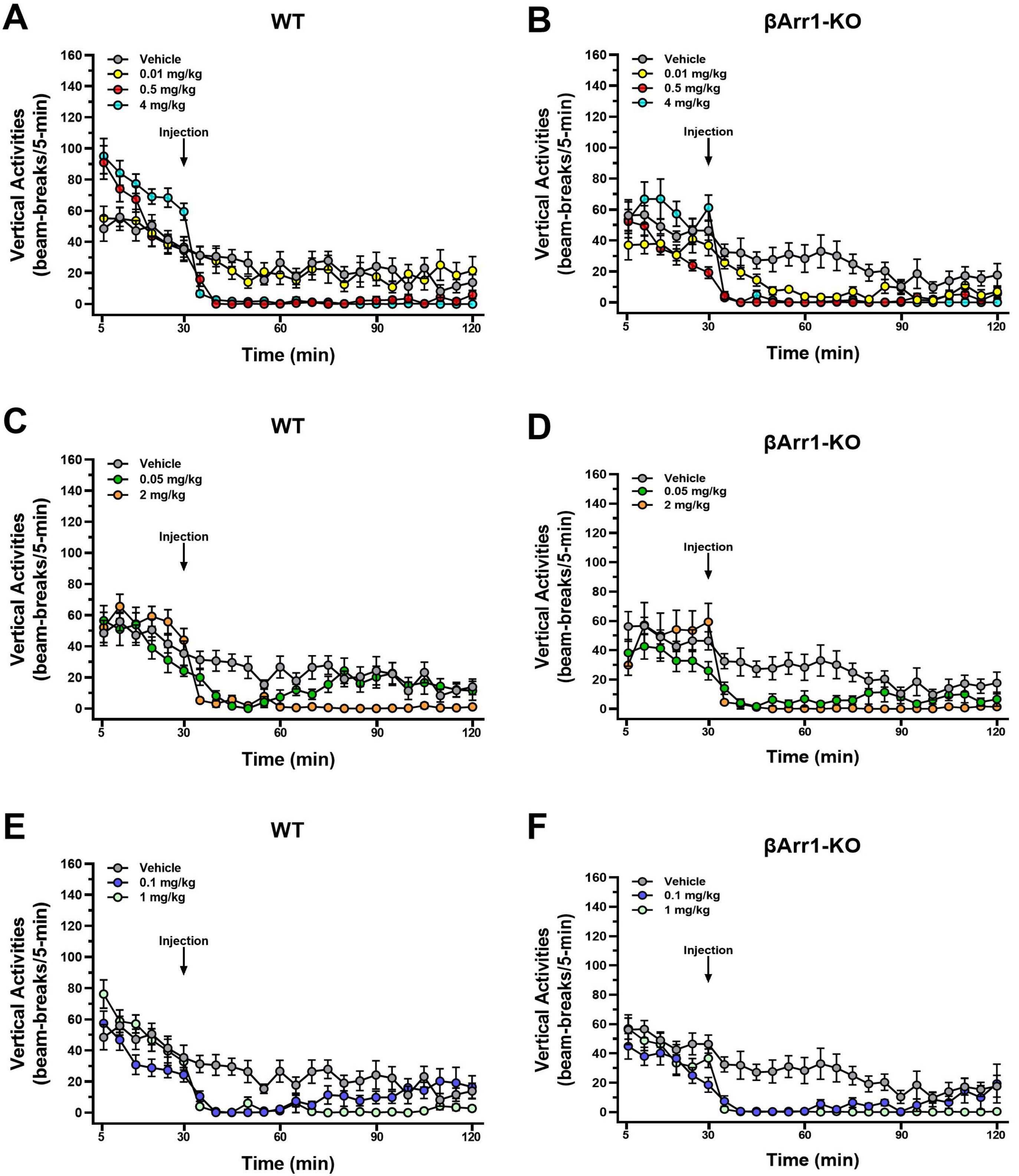
A
WT
B
βArr1-KO
C
WT
D
βArr1-KO
E
WT
F
βArr1-KO
Vehicle
0.01 mg/kg
0.5 mg/kg
4 mg/kg
0.05 mg/kg
2 mg/kg
0.1 mg/kg
1 mg/kg
Injection
Vertical Activities
(beam-breaks/5-min)
Time (min)

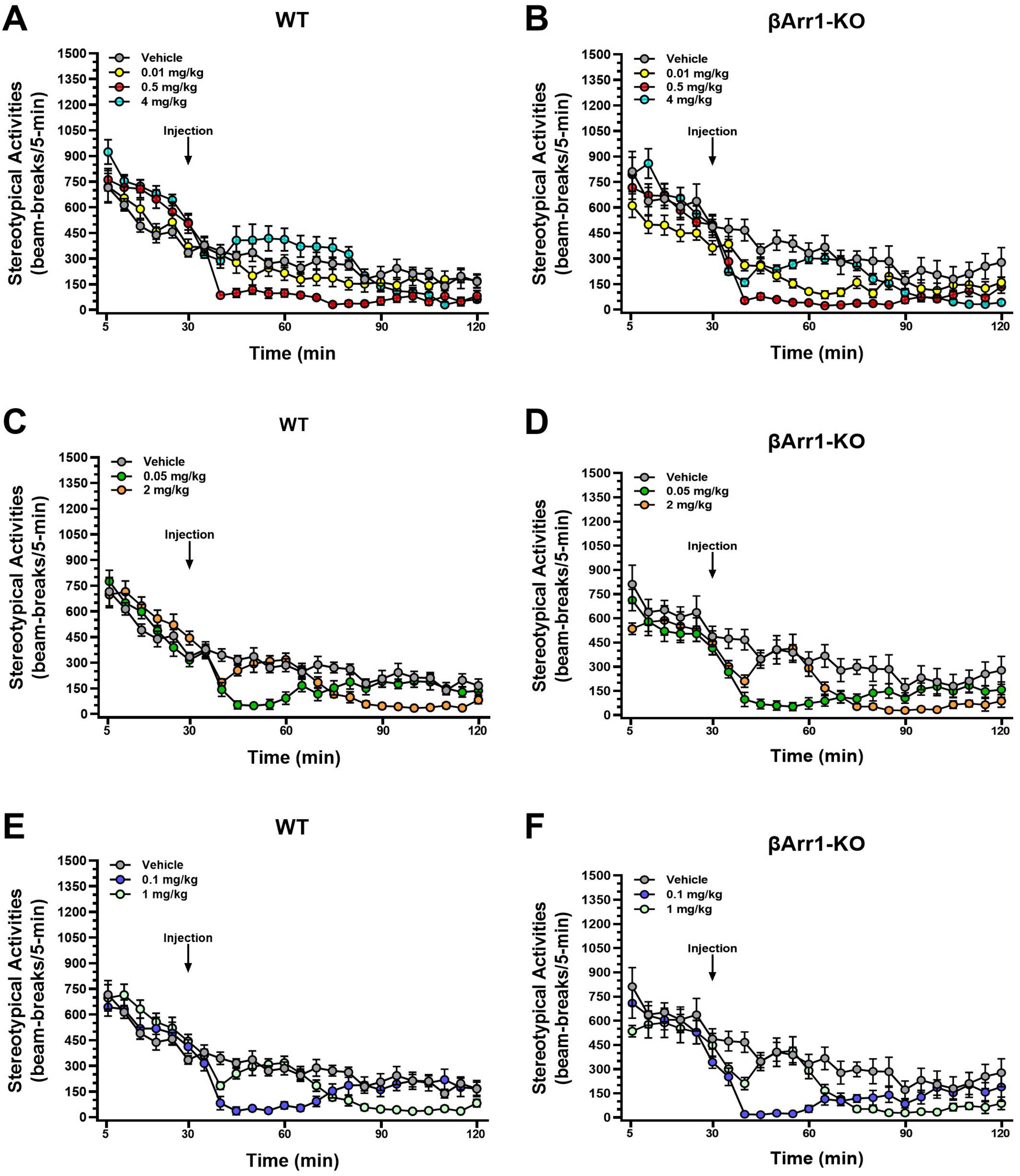
A
WT
B
βArr1-KO
C
WT
D
βArr1-KO
E
WT
F
βArr1-KO
Vehicle
0.01 mg/kg
0.5 mg/kg
4 mg/kg
0.05 mg/kg
2 mg/kg
0.1 mg/kg
1 mg/kg
Injection
Stereotypical Activities (beam-breaks/5-min)
Time (min)

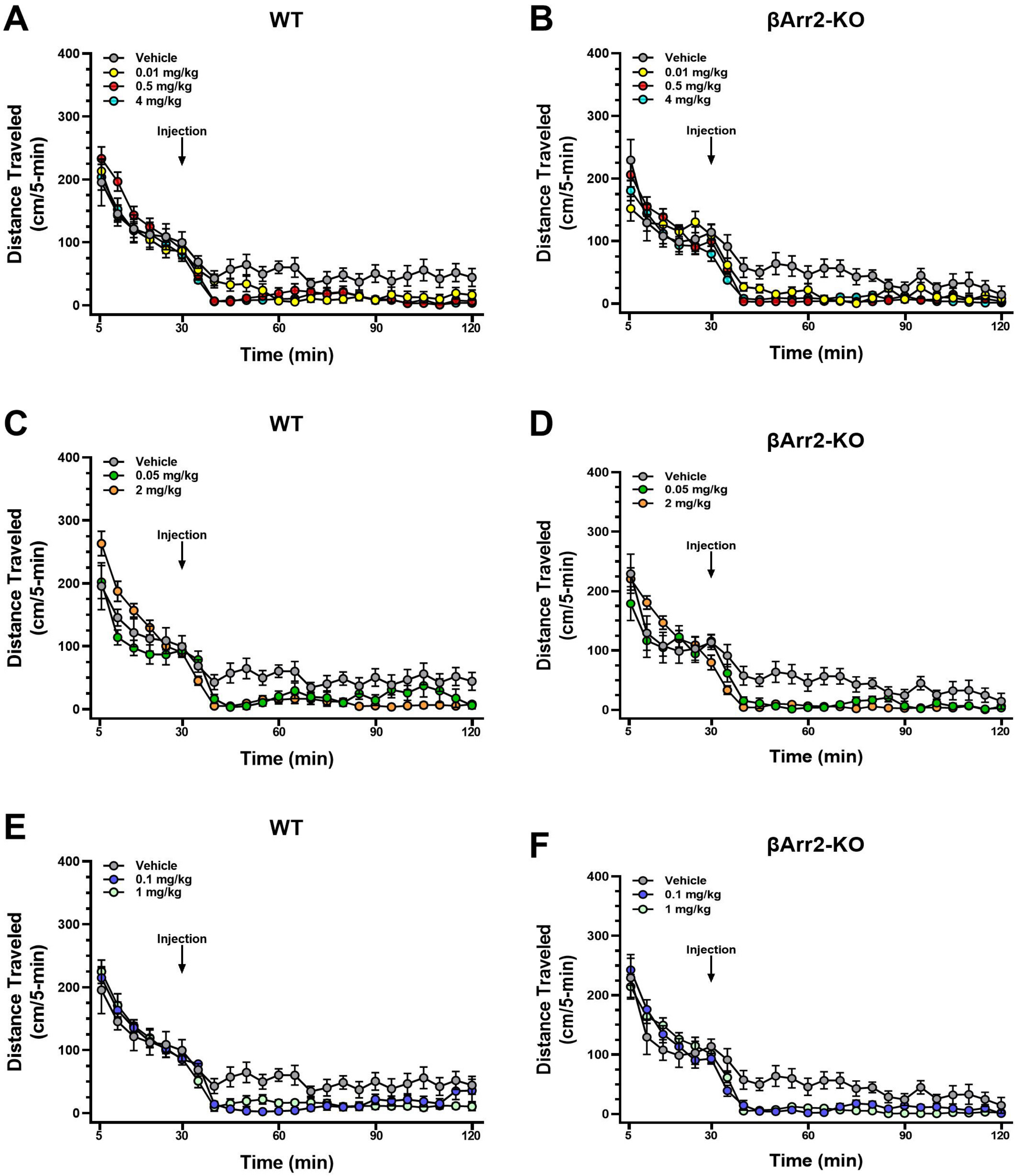
A
WT
Vehicle
0.01 mg/kg
0.5 mg/kg
4 mg/kg
Injection
Distance Traveled (cm/5-min)
Time (min)
B
βArr2-KO
Vehicle
0.01 mg/kg
0.5 mg/kg
4 mg/kg
Injection
Distance Traveled (cm/5-min)
Time (min)
C
WT
Vehicle
0.05 mg/kg
2 mg/kg
Injection
Distance Traveled (cm/5-min)
Time (min)
D
βArr2-KO
Vehicle
0.05 mg/kg
2 mg/kg
Injection
Distance Traveled (cm/5-min)
Time (min)
E
WT
Vehicle
0.1 mg/kg
1 mg/kg
Injection
Distance Traveled (cm/5-min)
Time (min)
F
βArr2-KO
Vehicle
0.1 mg/kg
1 mg/kg
Injection
Distance Traveled (cm/5-min)
Time (min)

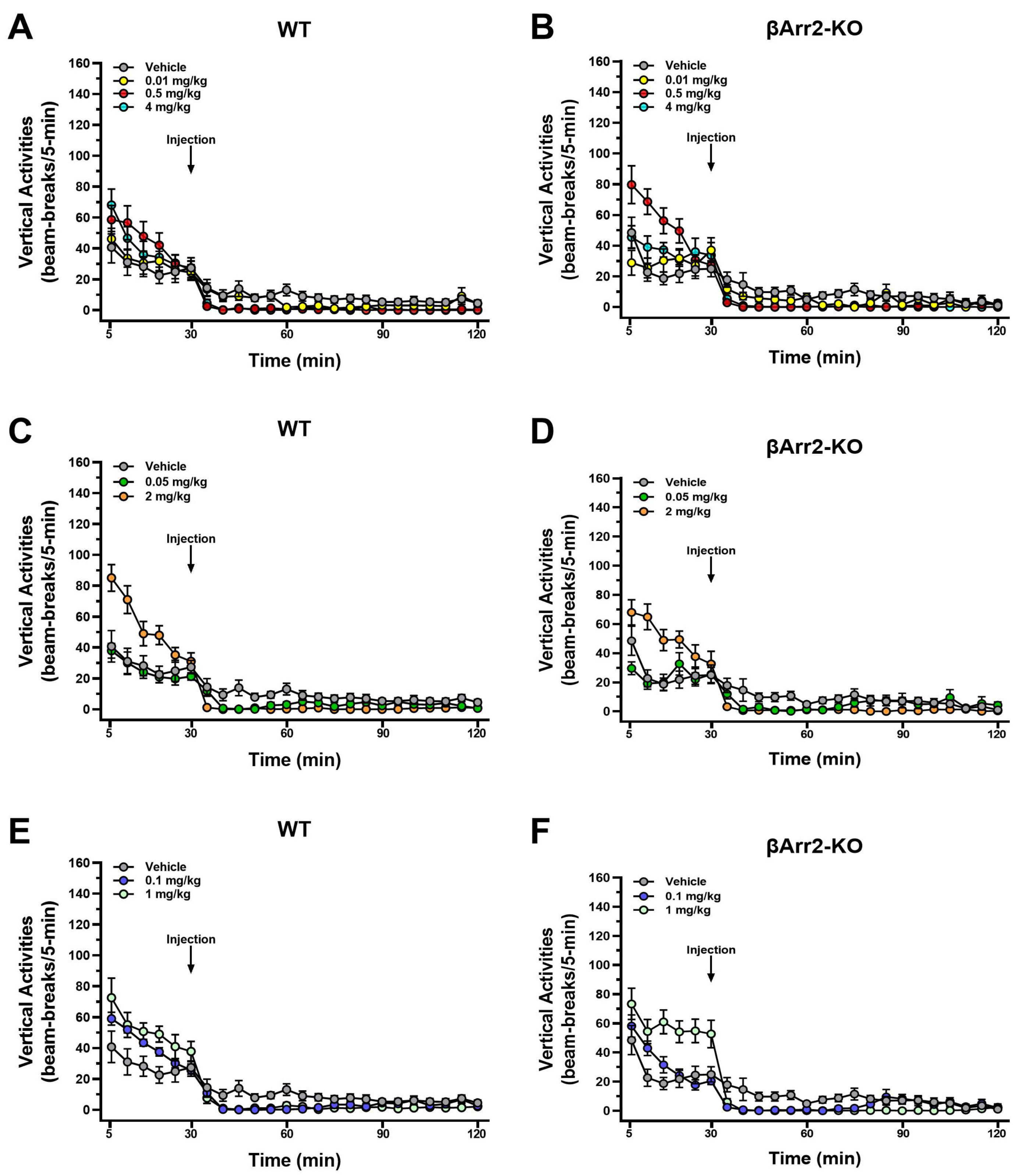
A
WT
B
βArr2-KO
C
WT
D
βArr2-KO
E
WT
F
βArr2-KO
Vehicle
0.01 mg/kg
0.5 mg/kg
4 mg/kg
0.05 mg/kg
2 mg/kg
0.1 mg/kg
1 mg/kg
Injection
Vertical Activities (beam-breaks/5-min)
Time (min)

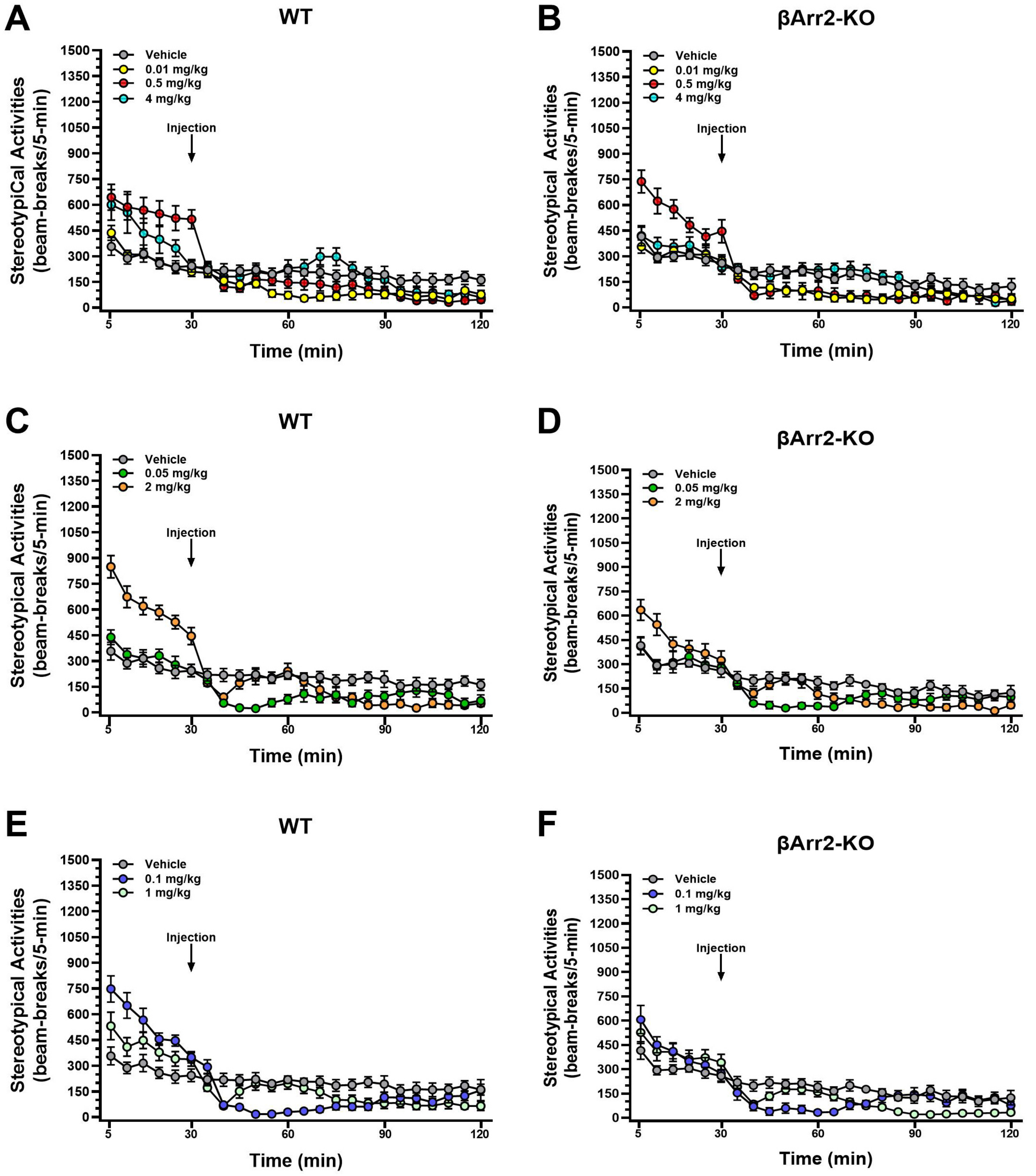
A
WT
B
βArr2-KO
C
WT
D
βArr2-KO
E
WT
F
βArr2-KO
Vehicle
0.01 mg/kg
0.5 mg/kg
4 mg/kg
0.05 mg/kg
2 mg/kg
0.1 mg/kg
1 mg/kg
Injection
StereotypiCal Activities (beam-breaks/5-min)
Stereotypical Activities (beam-breaks/5-min)
Time (min)

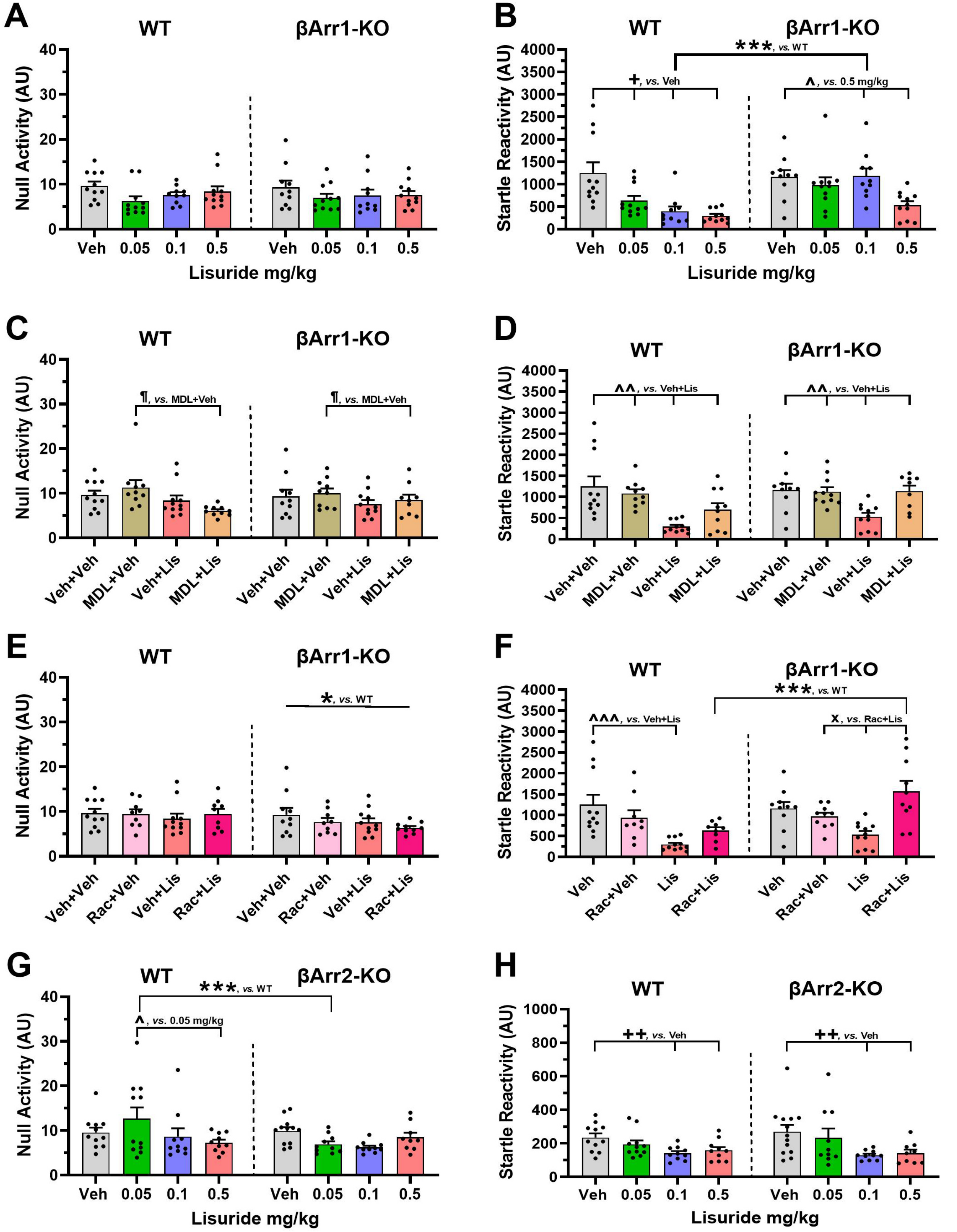

A
WT
βArr1-KO
Null Activity (AU)
Veh
0.05
0.1
0.5
Lisuride mg/kg
B
***, vs. WT
+, vs. Veh
^, vs. 0.5 mg/kg
Startle Reactivity (AU)
C
¶, vs. MDL+Veh
Veh+Veh
MDL+Veh
Veh+Lis
MDL+Lis
D
^^, vs. Veh+Lis
E
*, vs. WT
Rac+Veh
Rac+Lis
F
^^^, vs. Veh+Lis
x, vs. Rac+Lis
Lis
G
βArr2-KO
^, vs. 0.05 mg/kg
H
++, vs. Veh

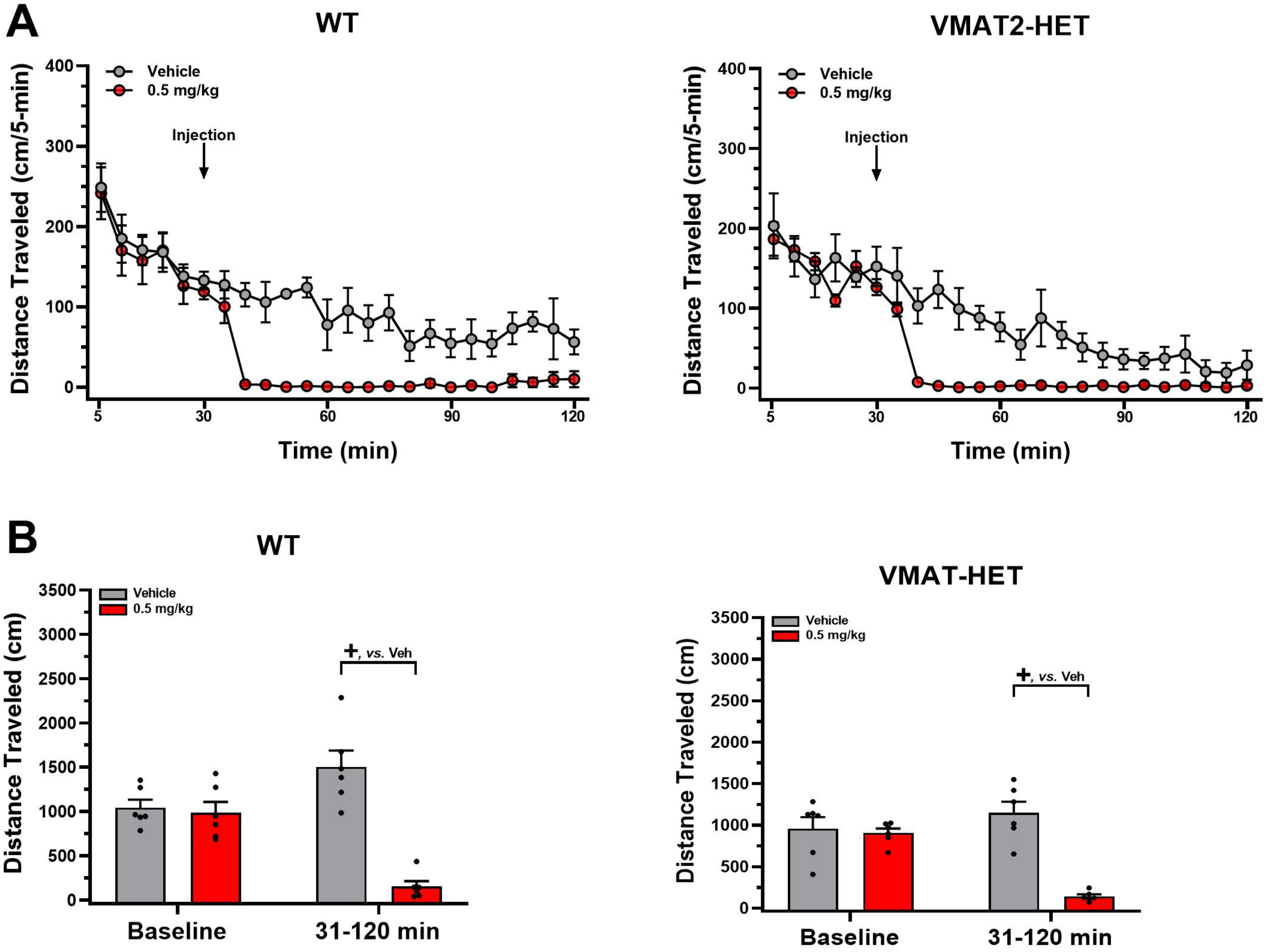
A
WT
VMAT2-HET
Vehicle
0.5 mg/kg
Injection
Distance Traveled (cm/5-min)
Time (min)
B
WT
VMAT-HET
Vehicle
0.5 mg/kg
+, vs. Veh
Distance Traveled (cm)
Baseline
31-120 min
